# Development of a Toolkit for High-Efficiency, Markerless, and Iterative Genome Editing in *Shouchella clausii*

**DOI:** 10.1101/2025.09.09.675032

**Authors:** Claudia Cappella, Carsten Jers, Lorenzo Ninivaggi, Maurizio Bettiga, Ivan Mijakovic, Gennaro Agrimi, Pasquale Scarcia

**Affiliations:** Department of Biosciences, Biotechnologies and Environment, University of Bari “Aldo Moro”, Campus Universitario, Via Orabona 4, 70125, Bari, Italy; Novo Nordisk Foundation Center for Biosustainability, Technical University of Denmark, Kongens Lyngby, Denmark; Italbiotec Srl Società Benefit, Piazza della Trivulziana 4A, 20126, Milan, Italy; Systems and Synthetic Biology, Chalmers University of Technology, Gothenburg, Sweden

**Keywords:** *Shouchella clausii*, *Bacillus clausii*, Electroporation, Markerless genome editing, Temperature-sensitive replication, Gram-positive bacteria, Homologous recombination

## Abstract

*Shouchella clausii* is a spore-forming, Gram-positive bacterium with intrinsic antibiotic resistance and promising potential in biotherapeutics, industrial biotechnology, and environmental applications. Its genetic intractability—due to a rigid cell wall and lack of natural competence—has limited its development as a microbial chassis. To facilitate its genetic transformation, a hyperosmotic electroporation protocol was optimized using cell wall weakening agents, achieving efficiencies comparable to other recalcitrant bacilli. A comprehensive and reusable genetic toolkit was developed centred on a temperature-sensitive *E. coli–S. clausii* shuttle vector (pM4B522), specifically engineered for compatibility with Golden Gate assembly. The plasmid backbone includes a spectinomycin resistance marker and an integrated red fluorescent protein reporter for transformants selection. A removable AmilCP chromoprotein cassette streamlines the assembly process by enabling blue/white screening in *E. coli*. This platform has demonstrated its versatility in genome editing for both *S. clausii* and *Bacillus subtilis*, as evidenced by its use in several applications: (i) sequential, markerless deletions of the non-essential catabolic genes *xylA* and *lacA* in *S. clausii* DSM 8716, with a success rate exceeding 60%; (ii) replacement of the *lacA* coding sequence with a GFP coding sequence, resulting in fluorescence induction in lactose-supplemented medium; (iii) introduction of a single-base substitution generating a premature stop codon in *lacA*, showcasing scar-free point mutagenesis; and (iv) transfer of the system to *B. subtilis* 168, highlighting its broader applicability across Gram-positive bacteria. Given the precision and scarless nature of these genetic modifications, this toolkit holds strong potential for the development of next-generation probiotics and synthetic biology applications.

## 1. Introduction

*Shouchella clausii* (formerly *Bacillus clausii*) (Joshi et al. 2021) is a spore-forming, alkaliphilic, Gram-positive bacterium that has a long history of safe use, as well as a growing number of applications in the fields of probiotics, industrial and environmental microbiology (Joshi et al. 2021; Elshaghabee et al. 2017; Nielsen et al. 1995). Beyond its probiotic role, *S. clausii* has been investigated for non-clinical uses: *B. clausii* strain T fully degrades (mineralizes) cephalosporin antibiotics (cefuroxime, cefotaxime, cefpirome) in wastewater (Kong et al. 2019); isolate I-52 secretes an alkaline protease that retains after 72 h >75% activity in 5% SDS and ∼110% in 10% H₂O₂, a property valued in the detergent industry (Joo et al. 2003); spores and their glycopolypeptide biosurfactant have been incorporated into silica sol-gel coatings to inhibit biofilm-driven corrosion of carbon steel (Purwasena et al. 2023); and the rhizobacterial strain B8 promotes biomass accumulation in *Brassica napus* and *Medicago sativa*, highlighting its potential as a biofertilizer (Oulebsir-Mohandkaci et al. 2021). Since *S. clausii* is already approved for human use, its application in industrial, agricultural, or waste-remediation contexts faces fewer regulatory and practical hurdles.

The species has been commercialized worldwide for over 60 years in high-spore-density, over-the-counter spore formulations such as Enterogermina® (Italy; a four-strain cocktail, SIN/O-C/T/N-R, 2–4 billion spores per vial) and Erceflora® (South-East Asia and Latin America; 2 billion spores per vial). Furthermore, clinical randomized trials showed that strain 088AE lowers both the incidence and severity of antibiotic-associated diarrhoea when co-administered with broad-spectrum antibacterials (Maity and Gupta 2021), and additional studies report benefits in paediatric upper respiratory tract infections and persistent diarrhoea (Sudha et al. 2019) (Madempudi et al. 2022) (Dang et al. 2024). Crucially, comparative genomics has identified chromosomally encoded resistance determinants - including *erm(34)*, *aadD2*, and the β-lactamase *bcl-1* – in DSM 8716 and commercial strains; these loci are not flanked by insertion sequences or other mobile elements, no conjugative plasmids have been detected and no study to date has demonstrated horizontal transfer in vivo (Bozdogan et al. 2003; Girlich et al. 2007). A 2022 narrative review synthesizing more than thirty years of clinical use, found no documented cases of in vivo horizontal gene transfer (Acosta-Rodríguez-Bueno et al. 2022). This evidence reframes *S. clausii*’ s intrinsic antibiotic resistance profile from a regulatory liability to a clinical advantage, thereby supporting its use as an antibiotic-compatible live biotherapeutic capable of preserving or restoring the microbiota during antimicrobial treatment.

*S. clausii* offers a unique combination of a long-standing history of human consumption, Qualified Presumption of Safety (QPS) status, and compatibility with antibiotics. These attributes position it as a lower-risk chassis for therapeutic development (Acosta-Rodríguez-Bueno et al. 2022) and a particularly promising candidate for next-generation probiotics (NGPs) and live biotherapeutic products (LBPs) (Chang et al. 2019)..

To our knowledge, no stably replicating plasmid or reproducible transformation workflow has yet been established for *S. clausii*. This process is complicated by the presence of thick, highly cross-linked peptidoglycan cell wall and the absence of key proteins required for natural transformation in Gram-positive bacteria (Ahmed et al. 2022; Kovács et al. 2009).

In this work, we overcame key obstacles to genetic manipulation of *S. clausii* DSM 8716 by (i) optimizing a high-osmolarity electroporation method to reduce the impact of spore wall resistance, and (ii) designing a modular *Escherichia coli*–*S. clausii* shuttle vector. This vector serves as a flexible, reusable, tool for scarless gene engineering including deletions, insertions and point mutations. The resulting system enables transformation efficiencies suitable for standard molecular biology applications and supports stable recombinant gene expression.

## 2. Materials and Methods

### 2.1. Bacterial Strains and Growth Conditions

For routine cloning and plasmid propagation, *E. coli* DH5α was cultivated in Luria-Bertani (LB) medium at 37°C, either in liquid culture with shaking or on LB agar plates. When appropriate, spectinomycin was added at a final concentration of 50 μg mL^-1^ for selection.

*S. clausii* DSM 8716, *B. subtilis* 168 and derived strains were propagated in LB medium at 30°C, 37°C, or 42°C as required. Spectinomycin was added at a final concentration of 100 μg mL^-1^ when needed.

### 2.2. Antibiotic Susceptibility Assays

Antibiotic susceptibility of *S. clausii* DSM 8716 was assessed by culturing the strain in both liquid and solid LB media supplemented with various antibiotics at concentrations in the range of 1-100 µg mL^-1^. For liquid assays, cells were inoculated into LB broth containing the desired antibiotic and incubated at 37°C for 24 hours with shaking; growth was monitored by measuring optical density at 600 nm (OD₆₀₀). For solid media assays, overnight cultures were plated onto LB agar containing the appropriate antibiotic and incubated at 37°C for 48 hours and growth was determined by visual inspection of colony formation. The resistance of the strain was determined by presence or absence of visible growth at each concentration.

### 2.3. Molecular cloning procedures

Genomic DNA from *S. clausii* DSM 8716 and *B. subtilis* 168 was extracted using the DNeasy UltraClean Microbial Kit (Qiagen). Plasmid DNA was routinely isolated from E. coli DH5α cultures using E.Z.N.A. plasmid DNA Mini Kit I (omega BIO-TEK), following the manufacturer’s protocols.

For vector constructions, DNA fragments were amplified using the primers listed in Table S1 and relevant template DNA. PCR amplifications were performed using PrimeSTAR GXL DNA Polymerase (Takara Bio); the sequences of all homology arms are provided in Table S2.

Overlap extension PCR was performed as described by Shevchuk et al. (Shevchuk 2004). When required, PCR products were purified using E.Z.N.A. Cycle Pure Kit (omega BIO-TEK) and quantified using Nanodrop 1000 Spectrophotometer (Thermo Scientific).

Relevant DNA fragments were assembled using either Gibson Assembly (NEBuilder HiFi DNA Assembly, New England Biolabs), or by Golden Gate Assembly (BsaI HiFi - New England Biolabs) and T4 DNA ligase (Thermo Scientific)), following the manufacturer’s protocols.

Plasmid DNA constructs were introduced into *E. coli* DH5α via chemical transformation using the Inoue method for preparing supercompetent cells (Green and Sambrook 2020). Transformed cells were then plated on LB agar containing spectinomycin and incubated overnight at 37°C. Transformants were verified by colony PCR or plasmid extraction, followed by restriction analysis, and Sanger sequencing (service provided by Eurofins Genomics).

### 2.4. Gram Positive Bacteria Transformation

Transformation of *S. clausii* DSM 8716 was performed by electroporation, following a modified protocol based on Zhang et al. (Zhang et al. 2015). Briefly, to prepare competent cells, overnight cultures were diluted 1:16 into fresh LBSP medium (LB + 0.5 M sorbitol + 0.05 M KH=PO= + 0.05 M K=HPO=, pH 7.2) and grown to an OD=== of 0.7–0.8. To enhance cell wall permeability, cultures were treated with cell wall-weakening agents (0.5% threonine, 0.05% Tween 80, and 0.6% glycine) for 1.5 hour at 37°C. Cells were harvested by centrifugation at 4°C and washed four times with ice-cold electroporation buffer (0.33 M trehalose, 0.5 M sorbitol, 0.5 M mannitol, 0.5 mM MgCl₂, 0.5 mM K₂HPO₄, and 0.5 mM KH₂PO₄; pH 7.2). The cell pellet was resuspended in the same buffer at a final volume equal to 1/60 of the initial culture volume. Electroporation was performed with a Gene Pulser II electroporator (Bio-Rad) in 0.1 cm gap cuvettes at 2.2 kV, 25 μF, and 200 Ω, using 60 ng of plasmid DNA and 100 ng of linear DNA per 50 μL of cells. After electroporation, cells were allowed to recover in LBMS (LB + 0.5 M sorbitol + 0.38 M mannitol) at 37°C for 5 hours (30°C, when required), then plated on selective LB agar containing spectinomycin (100 μg mL^-1^). Plates were incubated at 37°C (30°C, when required) for 36 hours to select for transformants.

Transformation of *B. subtilis* 168 was carried out using the natural competence protocol (Yasbin et al. 1975). Competent cells were incubated with 100 ng of plasmid DNA for 1.5 hours at 30°C and then plated on selective LB agar.

Following transformation of *B. subtilis* and *S. clausii*, colony PCR screening was performed using a protocol adapted from the iGEM Team Technion Israel (http://2015.igem.org/Team:Technion_Israel), with substantial modifications as described below. Individual colonies of *B. subtilis* or *S. clausii* were picked from agar plates and resuspended in 10 μL of weak Tris-HCl buffer pH 8.0. The suspension was incubated on ice for 10 minutes, then subjected to three heating cycles (two minutes at 700 Watt in a microwave followed by one minute at –20°C). After the final microwave step, samples were incubated on ice for an additional 15 minutes. Two microliters of the resulting lysate were used directly as PCR template. PCR reactions included an initial denaturation step at 95°C for 10 minutes to inactivate DNases released during cell lysis.

### 2.5. Gene editing with pM4B522 and its derivatives

Cells transformed with the pM4B522 plasmid or its derivatives were plated on LB agar containing spectinomycin (100 µg/mL) and incubated at 30 °C for 48 hours to allow replication of the temperature-sensitive plasmid. The resulting colonies exhibited uniform red fluorescence due to chromosomal expression of RFP. These colonies were subsequently re-streaked onto fresh spectinomycin plates and incubated overnight at 42□°C, a non-permissive temperature that inhibits plasmid replication. Colonies that remained red under these conditions were presumed to have integrated the plasmid into the chromosome.

Selected red colonies were then inoculated into antibiotic-free LB medium and cultured at 30 C overnight. To eliminate any remaining cells harbouring the replicative plasmid, cultures were shifted to 42 C in LB supplemented with spectinomycin for 5 hours. Finally, cells were plated on non-selective LB agar and incubated at 30 C for approximately 30 hours. This step facilitated a second recombination event, resulting in plasmid loss and the appearance of white colonies. Colonies that lost red fluorescence and failed to grow in the presence of spectinomycin were isolated and verified by PCR and sequencing to confirm successful genome editing and the absence of plasmid backbone sequences.

### 2.6. Phenotypic Validation of Recombinant Strains

To confirm the expected phenotypes of recombinant strains, functional assays were performed. For assessment of β-galactosidase activity, colonies were plated on LB agar supplemented with 200 μg mL^-1^ X-gal (5-bromo-4-chloro-3-indolyl-β-D-galactopyranoside) and incubated at 37°C overnight. Colony colour (blue or white) was assessed to distinguish wild-type and mutant phenotypes, respectively.

Growth on various carbon sources was assessed using spot assays on M9 minimal agar plates supplemented with 2% glucose, lactose, or xylose, as appropriate. Plates were incubated overnight at 37°C, and growth was evaluated by visual inspection.

For fluorescence reporter assays, recombinant strains were grown either in LB medium alone or in LB supplemented with 0.5% lactose for 24 hours at 37°C, to assess induction of the *lacA* promoter. Cultures were harvested, washed, and resuspended in sterile water to an OD₆₀₀ of 1. Fluorescence was measured using a PerkinElmer Luminescence Spectrometer LS 50B, with excitation/emission wavelengths of 558/583 nm for RFP and 488/510 nm for GFP. Results are expressed as mean ± standard deviation (SD) from three independent experiments; statistical significance was determined using one-way ANOVA with Tukey’s post-test, with P < 0.05 considered significant.

To assess α-amylase activity, recombinant *Bacillus subtilis* colonies were streaked onto LB agar plates supplemented with 10 g/L starch and incubated at 37 C for 24 hours. Following incubation, iodine crystals were placed in the empty half of the Petri dish, which was then sealed with the half containing the bacterial culture. This setup allowed iodine vapours to diffuse and react with the starch in the agar. Within minutes at room temperature, the starch turned dark blue, indicating its presence. Clear halos surrounding the wild-type colonies indicated starch hydrolysis due to α-amylase activity. Plates were photographed both before and after iodine exposure to document the results.

### 2.7. BLAST Search and Phylogenetic Analysis

*S. clausii* genomes available at NCBI were screened with the sequences of *B. subtilis* 168 XylA (BSU17600) and GanA (BSU34130) proteins using TBLASTN. The amino acid sequences were aligned with ClustalW and visualized with JALVIEW v2.11.4.1. The phylogenetic tree was generated from the ClustalW multiple-sequence alignment using the Maximum Likelihood method in MEGA11 and included the putative LacA sequences from *B. subtilis* and *S. clausii* strains.

## 3. Results

### 3.1. Transformation of *S. clausii* DSM 8716

To enable genetic engineering of *S. clausii* DSM 8716, we first sought to develop an efficient transformation protocol based on antibiotic selection. This required identifying an antibiotic to which the strain is sensitive, and for which a corresponding resistance marker was readily available. *S. clausii* DSM 8716 displayed resistance to several antibiotics commonly used as transformation markers, including erythromycin, chloramphenicol, and kanamycin but not spectinomycin (Table S3).

Given its sensitivity to spectinomycin and the availability of the corresponding resistance gene in our laboratory, for the scope of the present study, spectinomycin was selected for use as a marker in the transformation experiments.

To explore possible transformation strategies, the deletion of two non-essential genes—xylA, encoding xylose isomerase, and lacA, encoding β-galactosidase—was carried out using homologous recombination cassettes (Koo et al. 2017). These genes were selected due to the easily scorable phenotypes resulting from their loss in *Bacillus subtilis* (Gärtner et al. 1988; Daniel et al. 1997).

BLAST searches identified orthologues of XylA and GanA (LacA) of *B. subtilis* in *S. clausii* DSM 8716. For XylA, a single ortholog was found (BC8716 – 03450; 440 aa) shows 73.2% amino acidic identity to the *B. subtilis* enzyme (Figure S1).

For β-galactosidase, tBLASTP with the *B. subtilis* GanA protein returned two candidates: BC8716-10090 (AST96274.1; 682 aa; 53.9% identity) and BC8716-21430 (AST98362.1; 689 aa; 39% identity) (Figure S2). Across five additional *S. clausii* genomes (including the industrial strain KSM-K16; RefSeq NC_006582.1), orthologs of both loci were detected (Figure S2). Phylogenetic analysis (Figure S3) groups AST96274.1 with *B. subtilis* GanA and with the higher-identity *S. clausii* orthologs (96.6–100% identity among themselves), whereas AST98362.1 is weakly related and forms a separate clade. Therefore AST96274.1 was identified as the most likely GanA ortholog. Accordingly, a deletion was introduced at the *S. clausii lacA* locus (locus tag BC8716-10090).

A deletion construct targeting the *lacA* gene was designed to enable in addition, the insertion of *rfp* and *spec^r^* within its sequence (Figure 1A). To obtain a reusable source of the spectinomycin-resistance coding sequence, the *spec^r^* CDS was PCR-amplified from *B. subtilis* IHA01 (Härtl et al. 2001) and cloned into the shuttle vector pDR242a (Koo et al. 2017) obtaining the plasmid pM4B372.

**Figure 1.**
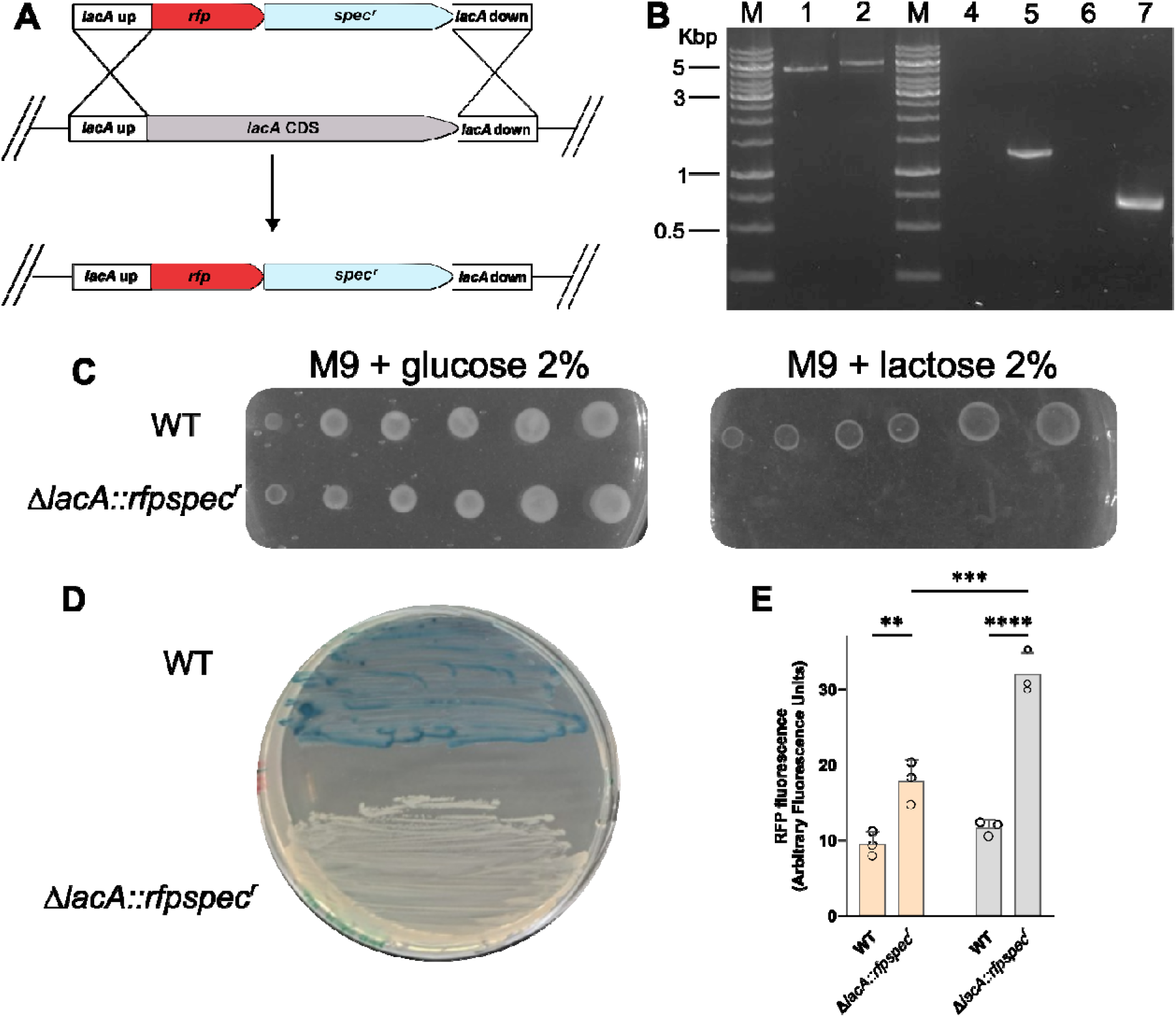
Replacement of *lacA* gene by homologous recombination. **A. Strategy for targeted replacement of the *lacA* gene in *S. clausii* DSM 8716 by homologous recombination.** B. Agarose gel analysis of colony PCR confirming cassette integration in *S. clausii* DSM 8716. Lanes M: DNA ladder (GeneRuler DNA ladder mix; Thermo Fisher); lanes 1-2: WT and Δ*lacA::rfpspec^r^* with external primers; lanes 4-5 -6 – 7: WT and Δ*lacA::rfpspec^r^* with internal *spec^r^* and *rfp* primers, respectively. **C. Carbon source utilization assay.** Serial dilutions of WT and Δ*lacA::rfpspec^r^*cultures spotted onto M9 minimal agar supplemented with 2% glucose or 2% lactose. **D. X-gal plate assay**. WT and Δ*lacA::rfpspec^r^* cells were streaked on LB agar supplemented with 200 μg mL^-1^ X-gal **E. RFP expression** Δ***lacA::rfpspec^r^* grown in LB or LB + lactose**. Fluorescence intensity of RFP expressed in Δ*lacA::rfpspec^r^* cells grown in LB (Light orange bars) and in LB supplemented with 0.5% lactose (Light gray bars). Bars represent mean ± SD (n = 3) of fluorescence Statistical analysis was performed using one-way ANOVA with Tukey’s post-test (\*\**P*< 0.01; \*\*\**P*< 0.001; \*\*\*\**P*< 0.0001).

For *lacA* integration cassette, ∼1 kb homology arms flanking *lacA* (Wu et al. 2019) were amplified from *S. clausii* DSM 8716 genome, while a *rfp* reporter gene was amplified from plasmid pBs1C-RFP (Radeck et al. 2013), and the *spec^r^* marker from pM4B372. These four fragments were assembled by overlap extension PCR (Shevchuk 2004) to generate a linear cassette designed for targeted integration into the *lacA* locus of the *S. clausii* genome (Figure 1A).

To introduce the *lacA* deletion cassette into *S. clausii* DSM 8716, we explored several transformation protocols. Notably, Kovács et al. (Kovács et al. 2009) reported that although some genes involved in natural competence are present in *S. clausii* KSM-K16—a strain closely related to DSM 8716—the absence of the regulatory gene *comK* likely renders the DNA uptake machinery inactive or highly inefficient. Consistent with these findings, natural competence protocols commonly used for *B. subtilis* (Yasbin et al. 1975) were tested but proved ineffective in *S. clausii* DSM 8716. Given these limitations, we adopted an electroporation-based approach instead. Starting from the high-osmolarity workflow of Zhang et al. (Zhang et al. 2015), four parameters, trehalose concentration, wall-weakening duration, DNA input, and recovery conditions, were optimized. The optimal set (0.33 M trehalose, 1.5 h weakening, 60 ng plasmid or 100 ng linear DNA per 50 µL cells, 5 h recovery) yielded transformation efficiencies of 7±3×10³ CFU µg⁻¹ of linear DNA —sufficient for routine genome engineering in *S. clausii* DSM 8716.

Successful integration of the rfp-spec^r^ cassette at the *lacA* locus was confirmed by colony PCR using three primer pairs. An external primer set flanking the *lacA* region yielded a 5.6 kb amplicon in recombinant clones, distinct from the 5.0 kb product observed in the wild-type strain. Only recombinant colonies produced PCR bands with the *rfp*- and *spec^r^*-specific primers, while wild-type controls showed no amplification with these internal primer sets (Figure 1B), confirming exclusive presence of both markers in engineered strains. Of the 15 colonies screened, 8 (54%) displayed the expected amplification pattern, indicating correct cassette integration. The ∼5.6 kb product obtained with external primers, spanning both homology-arm junctions, was gel-purified and subjected to Sanger sequencing, which verified precise integration and sequence fidelity at the insertion site. Notably, recombinant colonies were not visibly red on plates, consistent with the use of a promoter less *rfp* (reporter detectable only by PCR/fluorescence imaging).

Disruption of *lacA* was assessed using two phenotypic assays. Growth was evaluated on M9 minimal medium supplemented with either 2% glucose or 2% lactose as the sole carbon source (Figure 1C). Both WT and Δ*lacA::rfpspec^r^*strains grew on glucose, but only the wild type formed colonies on lactose, consistent with the inability of the Δ*lacA* mutant to catabolize this sugar. In parallel, wild-type and recombinant colonies were streaked on LB agar containing 200 µg mL^-1^ X-gal (Figure 1D). Wild-type colonies appeared blue, indicating functional β-galactosidase activity, whereas recombinant colonies were white, confirming loss of the *lacA* gene.

The phenotypic analysis of the Δ*lacA* strain suggests that the *lacA* gene (and hence not AST98362.1) encodes for the main beta-galactosidase enzyme in the *S. clausii* strain DSM8716, and it is essential for lactose metabolism.

Since the coding sequence of *lacA* was replaced also by the *rfp* gene and it could be used as a probe of the gene transcription level. To assess the heterologous expression of RFP, wild⍰type and recombinant strains were cultivated in LB, with or without 0.5% lactose, for 24 h, and their fluorescence was measured. As shown in Figure 1E, the recombinant strain exhibited a modest but statistically significant increase in RFP signal compared to the wild type grown in LB alone (p< 0.05). Notably, RFP fluorescence approximately doubled in the presence of lactose. These results suggest that *lacA* transcription is inducible by lactose, although a comprehensive characterization of the regulatory elements within the locus remains to be performed.

### 3.2 Development of a temperature-sensitive vector for S. clausii genetic modification

Having first established an efficient transformation protocol for *S. clausii* DSM 8716, we next engineered a versatile, recyclable, markerless editing vector. The resulting construct, pM4B522, was assembled by Gibson Assembly from two synthetic DNA fragments and comprises (Figure 2A): (i) the temperature-sensitive *repA^ts^* a replication origin from pE194 – obtained from the pMAD vector – which support replication in Gram-positive hosts at 30°C but is lost at 42°C (Arnaud et al. 2004); (ii) the ColE1/pM4B1 origin for high-copy replication in *E. coli*; (iii) a spectinomycin-resistance (*spec^r^*) cassette from pM4B372 (developed in this study) for selection in both *E. coli* and *S. clausii*; (iv) an *rfp* gene from pBs1C-RFP (Radeck et al. 2013) included as a visual marker to assess transformation in Gram-positive hosts; (v) an AmilCP chromoprotein cassette from plasmid JME5599 (Vidal et al. 2023) flanked by BsaI and BsmBI type IIS restriction sites, which is replaced during Golden Gate cloning and enables blue/white screening in *E. coli*.

**Figure 2.**
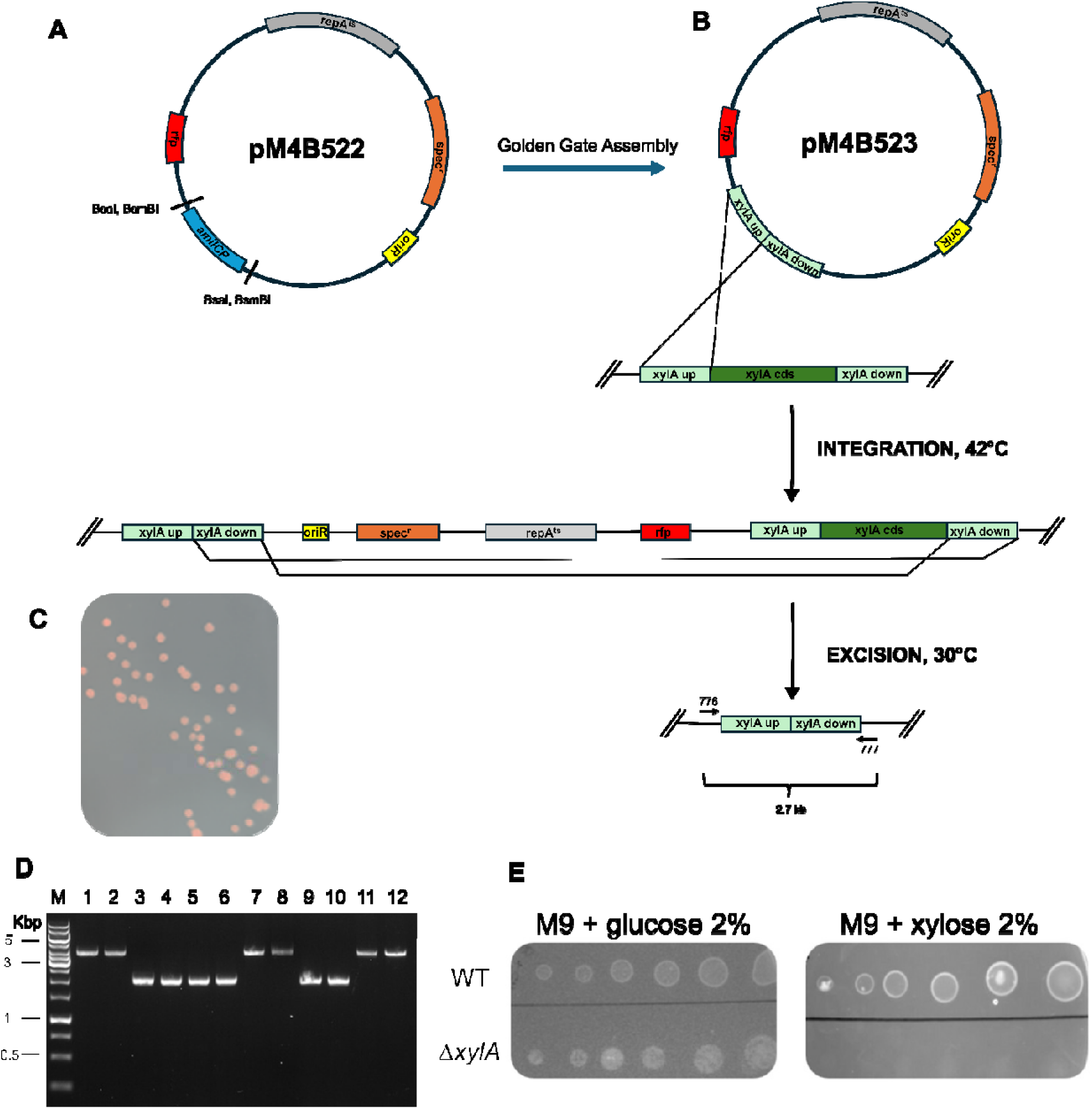
**A. Map of the pM4B522 plasmid.** **B. Two-step, markerless deletion of *xylA* in *S. clausii* DSM 8716 using the temperature-sensitive shuttle vector pM4B523**. The plasmid carries an *E. coli* replication origin (*oriR*, yellow), spectinomycin resistance cassette (*spec^r^*, orange), replication initiator *repA^ts^* (grey), and an *rfp* reporter (red). Upstream and downstream 1 kb homology arms (light grey) flanking the chromosomal *xylA* coding sequence (green). (i) Integration step: a single crossover between either arm and the chromosome inserts the entire plasmid, yielding a merodiploid intermediate. (ii) Resolution step: the second crossover can occur on either arm, yielding either the desired Δ*xylA* allele or restoration of the wild-type allele. In both cases, curing resulted in spectinomycin-sensitive clones, which were distinguished through colony PCR. **C. Plate showing *S. clausii* red colonies** grown at 30°C on selective medium containing spectinomycin (100 µg mL^-1^) after transformation with the recombinant plasmid pM4B523. **D. Agarose gel electrophoresis of colony PCR products from *S. clausii* strains**. Lane M: DNA ladder; Lane 1: wild-type (WT) control; Lanes 2–12: recombinant Δ*xylA* clones. Lanes with the Δ*xylA*-size amplicon correspond to deletion mutants; lanes with the WT-size amplicon are revertants (wild type after resolution). **E. Carbon source utilization assay for the** Δ***xylA* mutant**. Serial dilutions (left to right) of wild-type (WT) and Δ*xylA* cultures were spotted onto M9 minimal agar supplemented with 2% glucose (left) or 2% xylose (right) and incubated for 24 h at 37°C.

At permissive temperature (30°C), pM4B522 replicates in both *E. coli* and Gram-positive hosts. At non-permissive temperature (42°C), replication in Gram-positives is inhibited, thereby promoting recombination events and facilitating markerless genome editing.

The intact pM4B522 backbone expresses the chromoprotein AmilCP, resulting in blue colonies in *E. coli*. During Golden Gate Assembly, the entire *amilCP* cassette—flanked by BsaI and BsmBI sites—is excised and replaced by two homology regions (and, if required, any additional DNA cargo) targeting the desired genomic locus. Disruption of AmilCP results in white *E. coli* colonies, allowing immediate visual screening for correctly assembled constructs prior to sequencing. The resulting plasmids retain temperature sensitivity, and the *rfp* and the *spec^r^* modules for selection in Gram-positive hosts. The size of pM4B522 was reduced to approximately 6.9□kb—significantly smaller than the ∼9□kb of other temperature-sensitive vectors (Arnaud et al. 2004), to enhance ease of manipulation and improve transformation efficiency. This streamlined architecture served as the common backbone for all subsequent genome edits such as gene deletions, insertions, and point mutations described in the following sections.

The system is designed to allow scarless genome editing via double crossover, enabling precise modifications without the permanent introduction of antibiotic resistance markers. Such an approach increases the flexibility of strain engineering and permits multiple rounds of genome editing using the same selectable marker.

### 3.3. The pM4B522 plasmid enables efficient gene deletion

To delete the *xylA* coding sequence, two ∼1 kb homology arms flanking the *xylA* cds were amplified from *S. clausii* DSM 8716 genomic DNA and assembled into pM4B522 by Golden Gate, replacing the *amilCP* cassette (Figure 2B). Loss of blue color in *E. coli* identified correctly assembled plasmids (white colonies), which were verified by EcoRI digestion and Sanger sequencing (construct pM4B523; backbone size 7.1 kb including the homology arms).

Plasmid pM4B523 was introduced into *S. clausii* by electroporation using the optimized protocol developed in this study and previously described, and cells were plated on LB agar supplemented with spectinomycin (100 µg mL^-1^). Plates were incubated for 48 h at the permissive temperature of 30 °C, that allows plasmid replication. The resulting colonies (10^3^ colonies μg^-1^ plasmid DNA) were uniformly red due to chromosomal expression of RFP and were fully resistant to spectinomycin (Figure 2C).

To induce a first homologous recombination event (single crossover), red colonies were re-streaked on fresh spectinomycin plates and incubated overnight at 42°C—a non-permissive temperature at which the pM4B522 replicon cannot replicate extra chromosomally. Surviving colonies thus contained the plasmid integrated into the chromosome at either homology arm.

To permit excision of the plasmid backbone via a second homologous recombination event, clones with chromosomally integrated plasmid were transferred to antibiotic-free liquid LB and grown overnight at 30°C. To eliminate cells retaining the replicative plasmid, cultures were incubated in LB medium supplemented with spectinomycin at 42°C. Subsequently, cells were plated on non-selective LB and incubated at 30°C, where plasmid loss was indicated by the disappearance of both the red phenotype and spectinomycin resistance. Putative double-crossover colonies were first screened for loss of the pM4B523 backbone. After the final growth step at 30 °C, the white colonies were re-plated on LB+spectinomycin. All colonies failed to grow, indicating a loss of the plasmid⍰borne *spec^r^* cassette. A PCR with primers internal to the plasmid backbone produced a 1.0⍰kb band in the primary red transformants, but no product in any white, spectinomycin⍰sensitive colony (Data not shown). Colonies satisfying these three criteria were selected for locus-specific PCR and Sanger sequencing. It should be noted that the second recombination event can result in either restoration of the wild-type allele or incorporation of the desired mutation, depending on which homology arm is retained - ∼50:50 expected from random resolution of the second crossover. Colony PCR using primers positioned outside the homology arms yielded a 4.7 kb amplicon in the wild-type strain and a 2.7 kb amplicon in Δ*xylA* candidates, consistent with the expected size reduction following deletion of the *xylA* coding region (Figure 2D). Out of eleven clones tested, six (55%) showed the 2.7 kb band and were classified as positive. These PCR products were gel-purified and subjected to Sanger sequencing across the junction, confirming a precise deletion of the *xylA* coding sequence with no residual scar.

On M9–xylose minimal medium (both solid and liquid), the Δ*xylA* strain exhibited no growth, while its growth on M9–glucose remained unaffected (Figure 2E). This indicates that the *xylA* gene is essential for xylose catabolism in *S. clausii* DSM8716, consistent with findings in *B. subtilis* (Gärtner et al. 1988).

To accelerate the process, we kept the original transformation conditions but shortened the curing timeline. After transformation, colonies were grown 48 h at 30 °C on spectinomycin. Plates were then replica-plated to the same medium and incubated overnight at 42 °C still with spectinomycin to select for integrants. The next day, cells were streaked on antibiotic-free medium and grown overnight at 30 °C to allow loss of the integration cassette and finally plated overnight at 37 °C without antibiotic for counter-selection of double-crossover mutants. This adjustment reduced the protocol by ∼1 day while preserving the efficiency and purity of double-crossover selection.

To demonstrate that pM4B522 can be reused for sequential genome edits, we decided to delete the *lacA* coding region in the Δ*xylA* background. A construct, pM4B551, was generated by replacing the AmilCP cassette with two ∼1 kb regions flanking the *lacA* coding sequence using Golden Gate assembly. Successful assembly was verified through blue/white screening in *E. coli* DH5α and confirmed by Sanger sequencing.

Following transformation of *S. clausii* with pM4B551 and application of the temperature-shift workflow described earlier, white, spectinomycin-sensitive colonies were screened using primers flanking the recombination site. The parental strain produced the expected 5.0 kb amplicon, while Δ*lacA* candidates (four out of five clones tested) yielded a 3.0 kb fragment (Figure 3A), consistent with the precise removal of the ∼2.0 kb coding *lacA* sequence. PCR products from two independent positive colonies were purified and Sanger-sequenced across both junctions, confirming a seamless, scar-free deletion.

**Figure 3.**
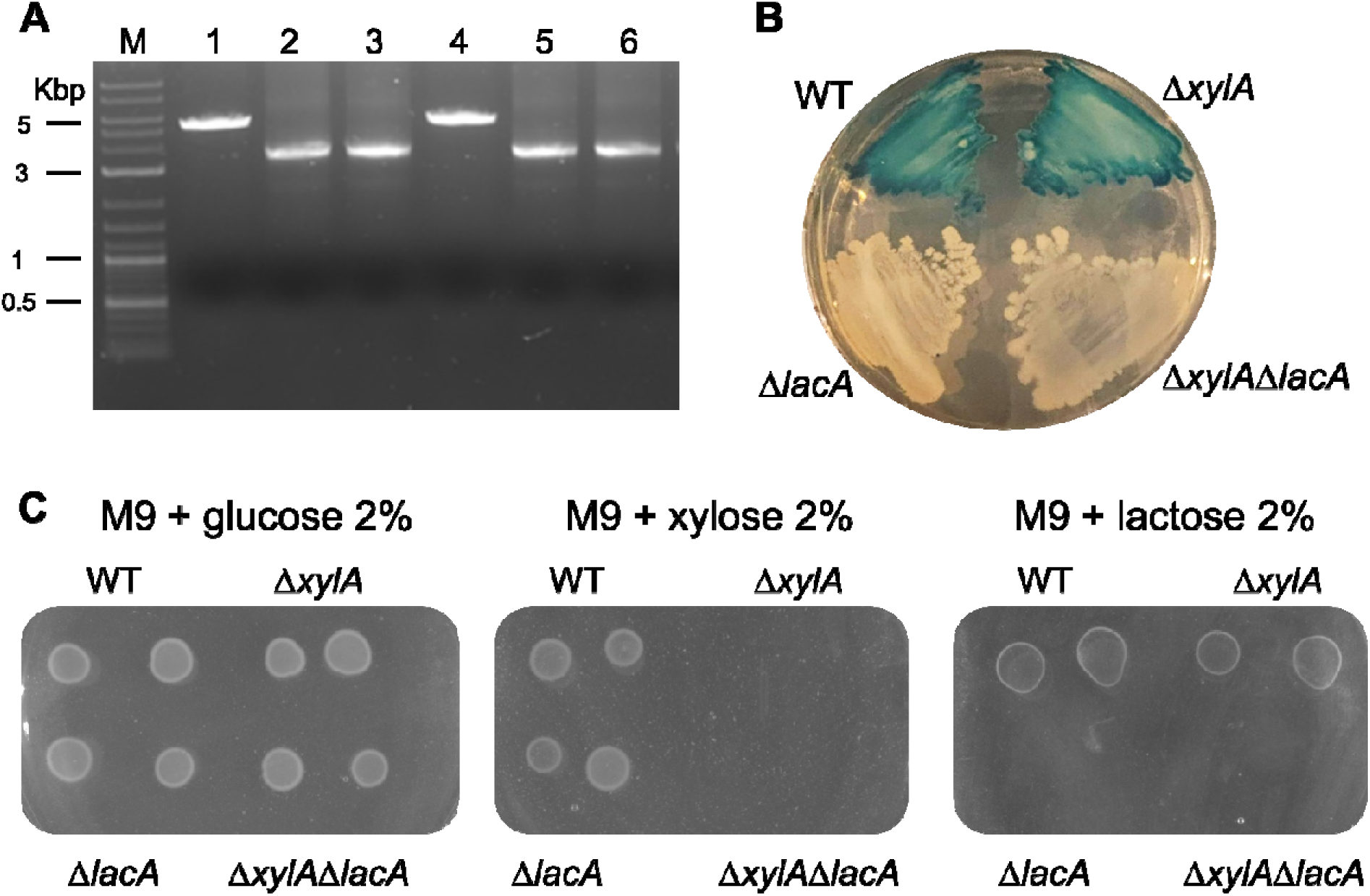
Molecular and phenotypic validation of *lacA* deletion. **A. Agarose gel electrophoresis of colony PCR products from WT and recombinant *S. clausii* strains**.1 Amplification with primers flanking the *lacA* locus yielded a ∼5 Kbp fragment in the wild-type strain (Lane 1) and a ∼3 Kbp fragment in recombinant clones with the *lacA* deletion (Lanes 2–6; with one negative clone in the lane 4). Lane M: DNA ladder (1 kb Plus Ladder, NEB). **B. X-gal LB agar plate.** *S. clausii* WT and mutant cells were plated on LB agar supplemented with X-gal 200 μg mL^-1^. The cells were distinguished by their ability to cleave the chromogenic substrate X-ga indicating the presence of a β-galactosidase activity. **C. Growth of *S. clausii* strains on M9 minimal medium supplemented with either glucose (left), xylose (centre), or lactose (right) as the sole carbon source**.

This result demonstrates that the markerless strategy can be applied iteratively: once the first edit is fixed, the same selectable cassette is available for subsequent rounds of editing without introducing additional resistance genes.

For comparison, the same *lacA* deletion was performed into the wild-type DSM 8716 background, yielding identical molecular and phenotypic results.

Phenotypic assays on LB-X-gal plates and on M9 minimal medium containing either glucose or lactose as the sole carbon source confirmed the genotype (Figure 3B - C).

### 3.4 pM4B522 can be used for insertion of heterologous genes

The *amilCP* cassette in pM4B522 was replaced with the *gfp* coding sequence (Wicke et al. 2017) flanked by the same 1 kb left and right homology arms of *lacA*, generating construct pM4B552. After transformation into *S. clausii* and completion of the temperature-shift curing protocol, white, spectinomycin-sensitive colonies were screened by colony PCR using external primers flanking the *lacA* recombination site. Recombinant clones produced the expected 3.7 kb amplicon, while the wild-type control yielded a 5.0 kb fragment, reflecting the ∼1.6 kb size difference between the *lacA* and *gfp* coding sequences. Of seven clones tested, six displayed the 3.7 kb band (Figure 4B). PCR products from three independent colonies were Sanger-sequenced, confirming seamless integration of *gfp* in place of the *lacA* coding sequence.

**Figure 4.**
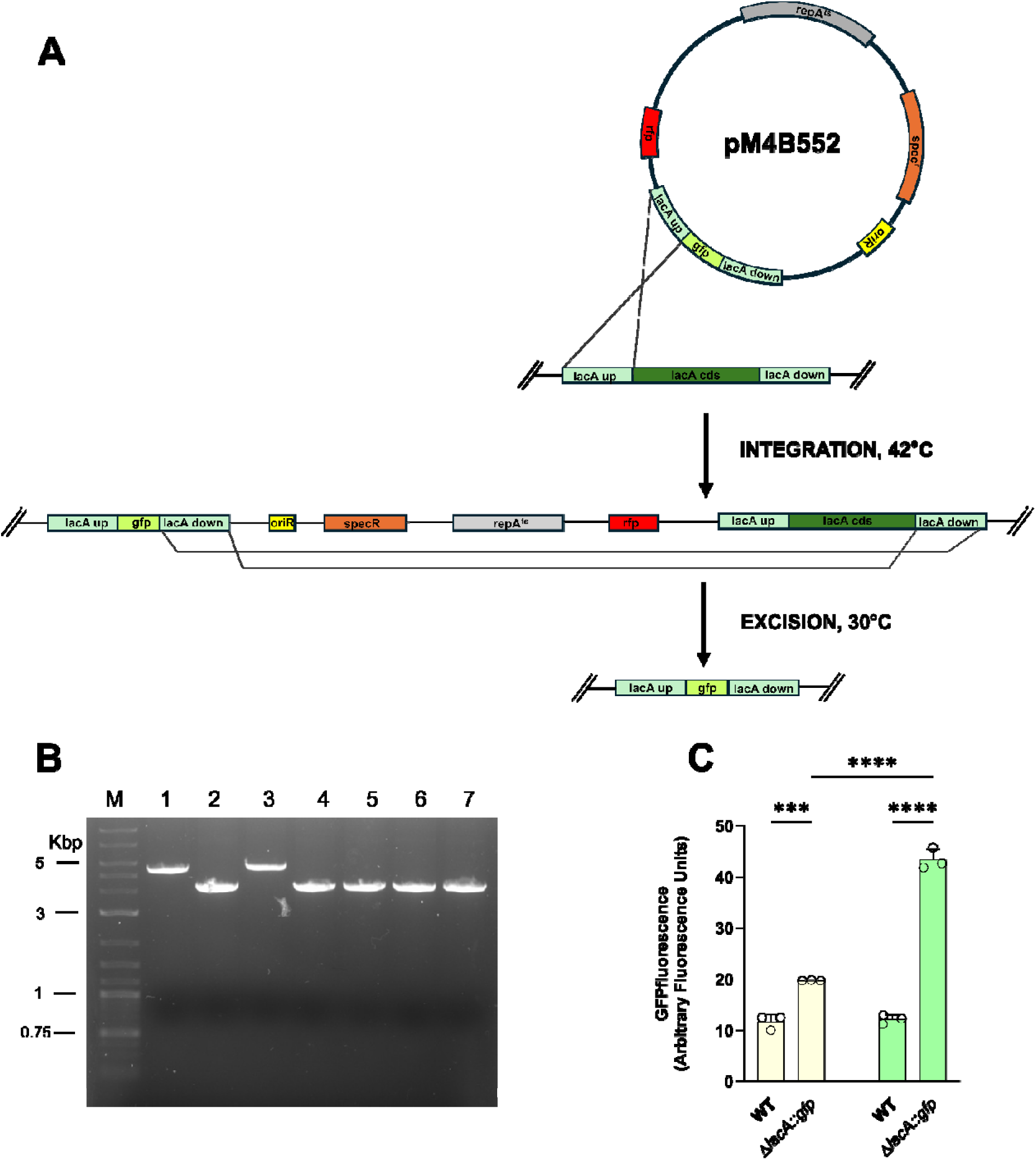
Replacement of *lacA* coding sequence with *gfp* in *S. clausii* DSM 8716. **A. Two-step, markerless replacement of *lacA* with the GFP coding sequence using the temperature-sensitive shuttle vector pM4B552**. The plasmid carries an *E. coli* replication origin (*oriR*, yellow), spectinomycin resistance cassette (*spec^r^*, orange), replication initiator *repA^ts^* (grey), and an *rfp* reporter (red). Upstream and downstream 1 kb homology arms (light green) flanking the chromosomal *lacA* coding sequence (green); in the pM4B552 plasmid, these homology arms flank the *GFP* coding sequence. (i) Integration step: a single crossover between either arm and the chromosome inserts the entire plasmid, yielding a merodiploid intermediate. (ii) Resolution step: a second recombination between the duplicated arms excises the vector backbone, leaving *GFP* at the locus. Black arrows indicate the direction of homologous recombination. **B. Agarose gel electrophoresis of colony PCR products from *S. clausii* strains**. Amplification with primers flanking the *lacA* locus yielded a ∼5.0 kb product in the wild-type strain (Lane 1) and a ∼3.7 kb fragment in recombinant clones containing the integrated *gfp* gene (Lanes 2–7), confirming successful replacement in lanes 2, 4, 5,6 and 7 and 6 (80% positive). Lane M: DNA ladder. **C. GFP fluorescence in *S. clausii* strains after 24 h growth under different conditions.** Fluorescence intensity of GFP expressed Δ*lacA::gfpspec^r^* cells grown in LB (Light yellow bars) and in LB supplemented with 0.5% lactose (Light green bars). Bars represent mean ± SD of three independent experiments. Statistical analysis was performed using one-way ANOVA with Tukey’s post-test (\*\*\**P*< 0.001; \*\*\*\**P*< 0.0001).

These findings were further supported by phenotypic assays (Figures 5C - D). Fluorimetric analysis showed significantly higher fluorescence in the recombinant strain compared to the wild type when grown in LB medium (p < 0.05). Moreover, fluorescence levels increased further when the recombinant clone was cultured in LB medium supplemented with 0.5% lactose (Figure 4C).

**Figure 5.**
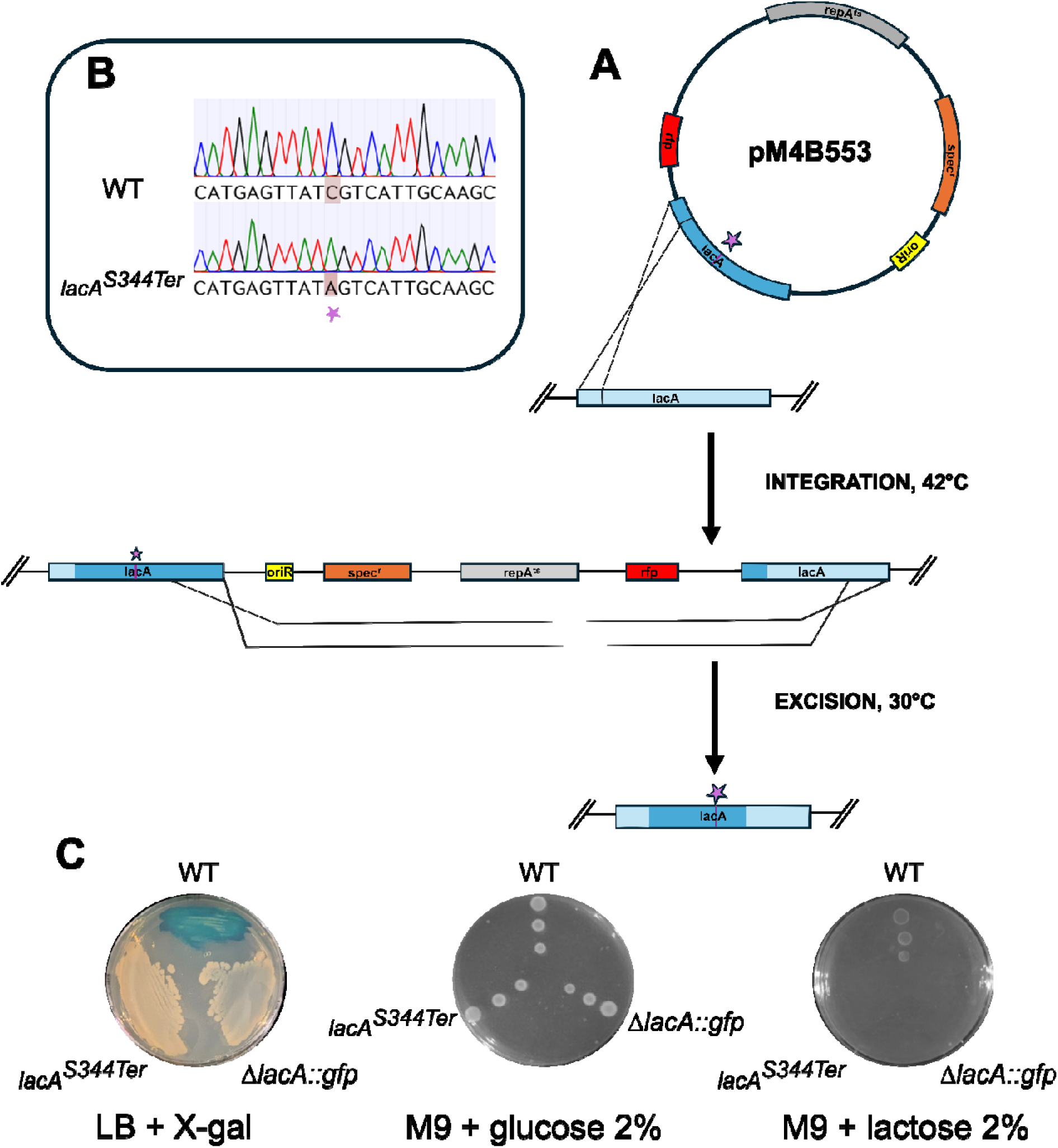
Allele replacement in the *lacA* coding sequence of *S. clausii* DSM 8716. **A. Two-step, markerless allele replacement in the *lacA* coding sequence using the temperature-sensitive shuttle vector pM4B553.** The plasmid carries an *E. coli* replication origin (*oriR*, yellow), spectinomycin resistance cassette (*spec^r^*, orange), replication initiator *repA^ts^* (grey), and an *rfp* reporter (red). (i) Integration step: a single crossover between either arm and the chromosome inserts the entire plasmid, yielding a merodiploid intermediate. (ii) Resolution step: a second recombination between the duplicated arms excises the vector backbone. The introduced point mutation is indicated by the purple star. Black arrows indicate the direction of homologous recombination events. **B. Sanger sequencing confirmation of the *lacA* C**→**A point mutation**. Electropherograms of the reverse strand for the parental strain (top) and the mutant clone (bottom). The highlighted region shows the single C→A transversion that converts the sense codon TCG (serine) into the in-frame stop codon TAG (stop codon), introducing a premature termination within the *lacA* coding sequence. **C. Phenotypic controls of** Δ***lacA::gfp* and *lacA^S344Ter^* recombinant clones. LB agar plate supplemented with X-gal showing colony color phenotypes**. Wild-type (WT) strain appears blue due to functional *lacA*, while *lacA^S344Ter^*mutant and Δ*lacA*::*gfp* mutants appear white, confirming replacement or inactivation of the *lacA* gene. **Growth on M9 minimal medium supplemented with glucose (left) or lactose (right).** Serial dilutions of WT, Δ*lacA*::*gfp*, and *lacA^S344Ter^* cultures were spotted onto M9 minimal agar and incubated for 24 h at 37°C.

### 3.5 The pM4B522 plasmid enables allele replacement

To evaluate the ability of pM4B522 to introduce single-nucleotide changes, we engineered a C→A transversion that converts codon 344 of *lacA* (TCG, encoding serine) into the stop codon TAG. The mutation was encoded directly in the primers used to amplify the two ∼2.2-kb fragments flanking the *lacA* coding sequence: the altered base was placed immediately downstream of the BsaI sites in both primers. After Golden Gate assembly, the mutagenized fragments replaced the wild-type segment in pM4B522, yielding the *lacA*-TAG allele (construct pM4B553). The construct was verified by Sanger sequencing before being used for transformation of *S. clausii*.

After the temperature⍰shift step, white, spectinomycin⍰sensitive colonies were screened by colony⍰PCR with the external *lacA* primers flanking the genomic locus; the amplicon size was identical to the wild type (5.1 kb), as expected. Sequencing of the PCR product revealed a single C-to-A substitution in eight of the twelve clones analyzed, leading to the generation of an in-frame stop codon (Figure 5B).

Phenotypic assays showed that the *lacA*^S344Ter^ mutants remained white on LB–X⍰gal plates and failed to grow when lactose was the sole carbon source, whereas growth on glucose was indistinguishable from the parental strain (Figure 5C).

These results show that pM4B522 backbone supports scar⍰free point⍰mutation editing and can be reused sequentially for deletions, insertions (expression of heterologous genes) and nucleotide substitutions.

### 3.6 Portability of the pM4B522 platform to *Bacillus subtilis* 168

To evaluate whether the marker⍰free system developed for *S.*□*clausii* could be transferred to other Gram⍰positive hosts, we tested it in *B.*□*subtilis*□168 using the chromosomal locus *amyE* as target.

Construct pM4B554—obtained by replacing the *amilCP* cassette with two ∼1.3 kb homology arms flanking the *amyE* coding sequence—was introduced into *B.*□*subtilis* by natural transformation. Following the temperature-shift curing step, white colonies sensitive to spectinomycin were screened via colony PCR. Using primers flanking the recombination region, a 2.8 kb amplicon— corresponding to the Δ*amyE* deletion—was obtained in five out of six colonies screened (80%), while the wild-type strain produced a 4.6 kb band (Figure 6A). Sanger sequencing across both junctions confirmed a scar-free deletion (data not shown). Loss of α⍰amylase activity was verified on starch plates (Figure□6C).

**Figure 6.**
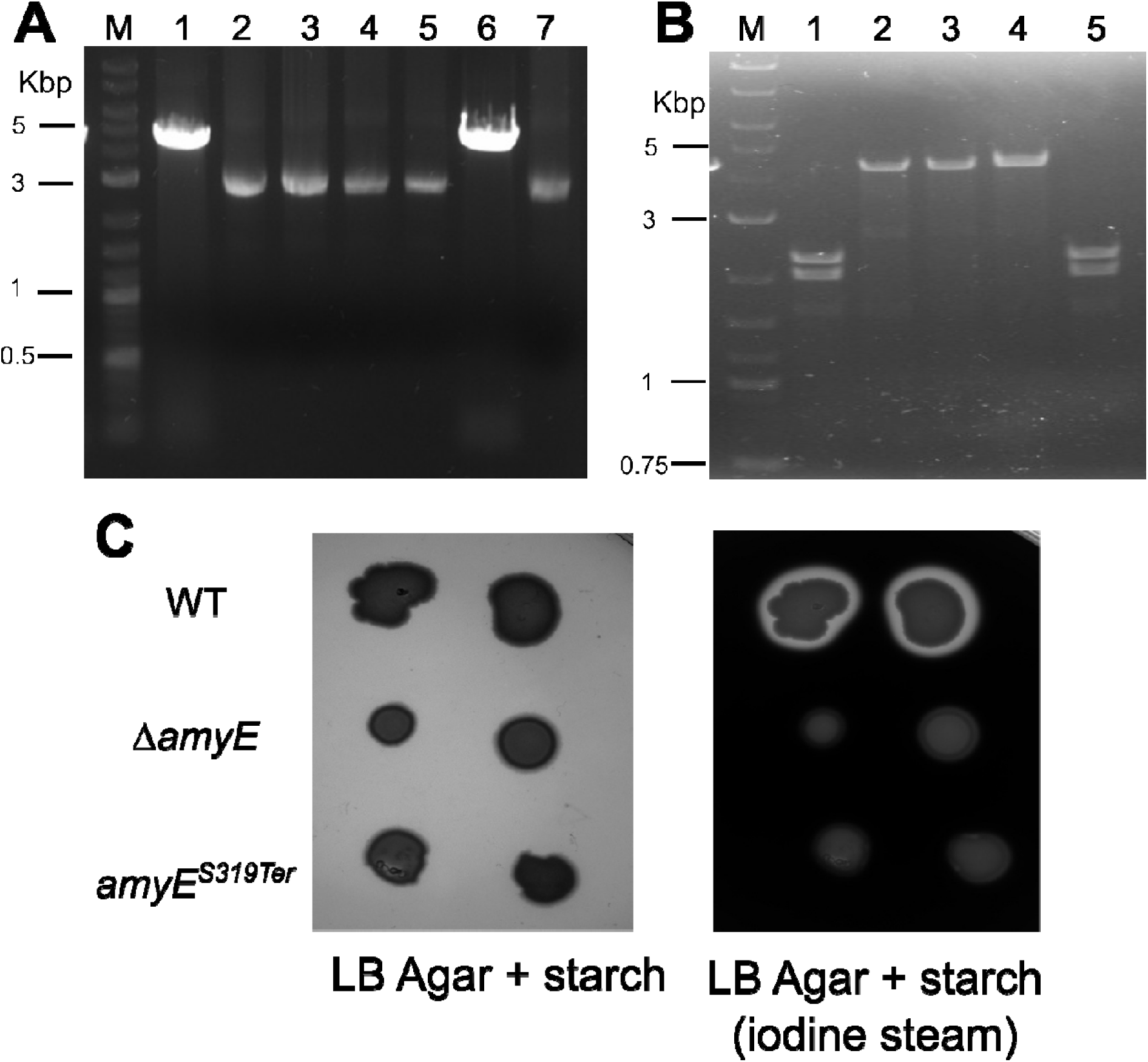
Markerless gene deletion and allelic replacement in *B. subtilis* 168. **A. Agarose gel electrophoresis of colony PCR products from *B. subtilis* strains** Δ***amyE* and *amyE^S319Ter^***. Lane M: DNA ladder; Lane 1: wild-type (WT) control; Lanes 2–7: recombinant Δ*amyE* clones. **B. Agarose gel electrophoresis of colony PCR products from *B. subtilis* strains digested with SalI.** Lane M: DNA ladder; Lane 1: WT control; Lanes 2–5: recombinant *amyE^S319Ter^* clones. Positive clones (lanes 2, 3, and 4) were not digested by SalI, while the wild type and a negative clone (lane 5) showed two bands after digestion. **C. Starch hydrolysis assay (*amyE* test) on LB agar plates supplemented with 1% starch.** Wild-type and mutant strains were streaked onto starch-containing LB plates. After overnight incubation, the plates were photographed (left panel) prior to exposure to iodine vapor (right panel), which interacts with starch. The presence of a clear halo surrounding colonies indicated starch degradation. Only the wild-type (WT) strain produced such a halo, confirming functional amylase activity. In contrast, neither the Δ*amyE* knockout strain nor the *amyE^S319Ter^* mutant showed halo formation, indicating loss of starch-degrading capability due to deletion or inactivation of the amyE gene.

A second construct, pM4B555, carried the entire *amyE* coding sequence but introduced a single C→A transversion at nucleotide +956 (codon 319), abolishing a SalI site (G<u>TCG</u>AC→G<u>TAG</u>AC) and generating an in⍰frame stop codon (<u>TAG</u>) in place of a Serine. The mutation was embedded in the overlapping region of the primers used to amplify the homology arms.

Following curing, colony PCR using primers flanking the recombination sites produced a 4.6 kb fragment, matching the wild-type length as expected. The absence of the SalI restriction site provided a quick screen for allelic replacement; three out of four clones tested (75%) resisted SalI digestion, indicating successful genetic modification (Figure 6B). Sequencing of PCR products from the three clones (data not shown) confirmed the presence of the intended single-base edit in all cases. Results were also confirming by phenotypic assay of amylase activity (Figure 6C).

These data demonstrate that pM4B522 supports both scar⍰free gene deletions and precise point⍰mutation editing also in *B. subtilis* 168 with high efficiency, mirroring the performance observed in *S. clausii*.

## 4. Discussion

The limited availability of genetic tools for *S. clausii* has long hindered its advancement—not only as a spore-forming chassis for biotherapeutic delivery, but also in broader industrial and environmental biotechnology applications. In this sense two major obstacles are (i) a highly cross-linked peptidoglycan layer that restricts DNA uptake (Donovan et al. 1987) and the absence of the competence master regulator ComK, which probably precludes an efficient natural transformation (Alcaraz et al. 2010) and (ii) the lack of a molecular toolkit for manipulation of its genome.

### 4.1 Transformation Protocol for *S. clausii* DSM 8716

By combining a trehalose–mannitol hyperosmotic buffer with mild cell-wall-weakening agents, we routinely achieved transformation efficiencies of approximately 7 × 10³ CFU/µg DNA—an order of magnitude higher than those previously reported for recalcitrant bacilli such as *Bacillus pumilus* (Danilova et al. 2022), and comparable to or exceeding those seen in *B. thuringiensis* (Lereclus et al. 1989). Although still below the ≥10□CFU µg⁻¹ reported for response-surface-optimized protocols in *B. subtilis* ZK (Zhang et al. 2015), these results represent, to our knowledge, the first reproducible protocol for transformation in *S. clausii* DSM 8716. Further studies are required to investigate natural competence in *S. clausii*. To this end, introducing regulatory elements such as the *comK* gene and the competence-stimulating peptide *ComS* from *B. subtilis*—which stabilizes the master regulator ComK—could help to evaluate their impact on natural competence in *S. clausii*.

### 4.2 Design and Implementation of a Recyclable, Marker-Free Genome Editing System

To overcome the absence of molecular tools for genome manipulation in *S. clausii*, a recyclable, marker-free allelic exchange vector was developed. This need led to the construction of pM4B522, a plasmid incorporating the replication origin from pMAD (Arnaud et al. 2004) – the pE194-derived *repA^ts^* module for temperature-sensitive replication in Gram-positive hosts – while redesigning all other functional modules. pM4B522 has an approximate size of 6.9 kb, making it significantly smaller than pMAD and other temperature-sensitive vectors like pKSV7 (Gao et al. 2017) and pKOR1 (Bae and Schneewind 2006). Additionally, the ∼1 kb AmilCP cassette is removed during Golden Gate assembly, reducing the final construct size to approximately 5.8 kb plus the homology arms—resulting in editing vectors that fall well within the 6–8 kb range known to yield optimal electroporation efficiency in *Bacillus*. Bidirectional BsaI/BsmBI landing pads facilitate seamless, scar-free cloning of inserts in a single-tube reaction. Screening is visual in both hosts: in *E. coli*, loss of amilCP produces white colonies, whereas in Gram-positive transformants the *rfp* reporter turns single⍰crossover colonies red making the curing step unambiguous. Once the plasmid has been excised following a second crossover event, no antibiotic marker remains in the chromosome; consequently, the same *spec^r^* backbone can be recycled indefinitely without changing antibiotics between cloning and transformation. Because pM4B522 retains the pE194⍰derived *repA^ts^* origin used by pMAD, it is plausibly portable to many low⍰GC Gram⍰positive hosts where pMAD replicates. In short, pM4B522 preserves the proven *repA^ts^* logic of pMAD but couples it to modern Type IIS modularity, a smaller electroporation payload, and dual⍰host colour screening—maintaining the key advantage of rapid temperature-sensitive plasmid loss without the need for counter⍰selection.

Using this toolkit to delete coding sequences or introduce stop codons, we confirmed that proteins *LacA* and *XylA* are essential for lactose and xylose metabolism in *S. clausii* DSM 8716, respectively. These proof-of-concept edits further emphasize the platform’s value for functional genetics research in *S. clausii*.

### 4.3 Comparison with Alternative Markerless Gene-Editing Technologies

Several strategies now enable scar-free genome editing in *Bacillus* spp. (i) CRISPR/Cas counter-selection: multiplex toolkits use a plasmid-borne *cas9* and a repair template to enrich for correctly edited cells, achieving efficiencies of 95% or higher (Altenbuchner 2016; Westbrook et al. 2016; Zou et al. 2022). While this approach is rapid, it carries the risk of off-target double-strand breaks or unintended gene silencing during extended Cas9 expression. These effects, driven by sgRNA-dependent binding of Cas9 or dCas9 to non-target DNA sequences, have been reported in various bacteria, including *Bacillus* (Rostain et al. 2023). (ii) Site-specific recombinase systems (Cre-loxP, Xer/dif, FLP-FRT) excise an antibiotic marker flanked by recognition sites, delivering seamless edits after a second induction step, yet they leave a 34- to 82-bp recombination “scar” (Yan et al. 2008). (iii) Counter-selectable metabolic markers—such as upp with 5-fluorouracil, mutant pheS with p-chlorophenylalanine, or codA with 5-fluorocytosine—enable fully markerless genome edits. However, they require host-specific optimization and often result in lower recombination efficiencies (Fabret et al. 2002). (iv) Recombineering using phage RecT can introduce point mutations without the need for selection; however, in Bacillus, the method is currently limited to specialized strains expressing heterologous annealases and still relies on PCR screening to identify the rare correctly edited clones (Sun et al. 2015).

By exploiting endogenous homologous recombination and a temperature-sensitive replicon instead, pM4B522 leaves behind no foreign nuclease genes, residual recombinase sites or acquired antibiotic markers. In general, no additional DNA sequence remains in the host’s genome except for the intended genetic modification once cured. This marker-free, scarless system aligns with the US FDA’s Chemistry, Manufacturing and Control guidance for live biotherapeutics, which strongly discourages the presence of acquired resistance genes. (FDA - *Early Clinical Trials with Live Biotherapeutic Products: Chemistry, Manufacturing, and Control Information; Guidance for Industry*).

### 4.4 Toward a lactose-responsive reporter in *S. clausii*

Having established reliable gene integration in *S. clausii*, the next requirement is access to efficient promoters and translation control. In *Bacillus* spp., synthetic promoter libraries enable predictable tuning of transcription (Okay 2024; Hammer et al. 2006). In *B. subtilis*, multiple inducible options exist (e.g., PxylA, Pspac, Pgrac), which could be ported as needed. Our chromosomal *lacA*→*rfp/gfp* replacement, although yielding modest absolute fluorescence, demonstrates that transcription at the *lacA* locus responds to lactose. This proof-of-concept provides a practical entry point to develop a lactose-inducible expression system in *S. clausii*, which can be rapidly implemented on our Golden-Gate–ready backbone.

### 4.5 Toward Next-Generation Live Biotherapeutics

From an application perspective, *S. clausii* spores withstand gastric acidity, germinate in the small intestine, and have an excellent human safety record (Acosta-Rodríguez-Bueno et al. 2022). Engineering this chassis opens the door to in situ delivery of bioactive molecules, from anti-inflammatory cytokines to metabolic hormones—an approach already validated in *Lactococcus lactis* (Steidler et al. 2003) and *E. coli* Nissle (Isabella et al. 2018). Since pM4B522 introduces only the intended genetic modification without leaving behind any foreign DNA, it significantly reduces both regulatory and technical hurdles for implementing protein-secretion strategies in an *S. clausii* background—leveraging the strain’s natural antibiotic compatibility alongside the genetic adaptability needed for next-generation live biotherapeutics.

## 5. Conclusions

In conclusion, the developed electroporation protocol combined with the recyclable Golden-Gate vector pM4B522 transforms the reference strain *S. clausii* DSM 8716 from a genetically recalcitrant organism into a tractable chassis for synthetic biology. By minimizing the plasmid backbone size, eliminating residual antibiotic markers, and avoiding the use of heterologous nucleases, this platform overcomes longstanding technical barriers while aligning with current FDA guidelines for live biotherapeutics—specifically, avoiding the acquisition of antimicrobial resistance markers and minimizing residual foreign DNA. Although regulatory classification ultimately depends on jurisdiction and product-specific risk assessment, our markerless workflow reduces typical regulatory burdens by eliminating resistance genes and ensuring editing plasmids are not retained. This combination marks a significant improvement over previous methods, offering reproducible transformation efficiencies in the range of 10³ CFU µg⁻¹ and supporting scar-free and iterative deletions, insertions and allele replacements. Since pM4B522 uses the pE194 *repA^ts^* origin, it is plausibly portable to other Gram⍰positive bacteria; future work will benchmark its performance beyond *S. clausii* and *B. subtilis*. In summary, the developed toolkit provides a robust platform for the genetic engineering of *S. clausii*, enabling a broad spectrum of applications—from the creation of next-generation probiotics and live biotherapeutic products (LBPs) to advancements in metabolic engineering, biocatalysis, bioremediation, and agricultural biotechnology.

## Conflinct of interest

The authors declare that there are no conflicts of interest associated with this manuscript. No financial, personal, or professional relationships influenced the work reported in this study.

## Availability of data and materials

The datasets generated or analyzed during this study are available from the corresponding author upon reasonable request.

