## Supplementary material for "Development of a Toolkit for High-Efficiency, Markerless, and Iterative Genome Editing in *Shouchella clausii*"

### Table S1. List of primers used in this work

| Name | Sequence | Use |
|---|---|---|
| 757 | aaagaattcttgacatttttcttgtggatctgtataataaagaataatta | Amplification of *specR* cassette – forward primer with EcoRI site |
| 758 | atactcgagatcgatcaatagttacaaattgtttcacta | Amplification of *specR* cassette – reverse primer with + XhoI site |
| 807 | agtttgcgtggcaaatgg | *lacA* upstream region – forward primer |
| 50 | acgtcttcggaggaagccacgccaagcgattgtcgt | *lacA* upstream region – reverse primer, overlaps with *rfp* |
| 51 | acgacaatcgcttggcgtggcttcctccgaagacgt | *rfp* – forward primers, overlaps with *lacA* up |
| 52 | ctctcctttctcgcctgcttaagcaccggtggagtgacg | *rfp* – reverse primer, overlaps with *specR* |
| 716 | gcaggcgagaaaggagaggagggaggaaaggcagga | *specR* – forward primer(1) |
| 717 | cgaggctcctgtcactgccgccgtatctgtgctctc | *specR* – reverse primer(1) |
| 809 | gcagtgacaggagcctcgatgtcattaatgactgctg | *lacA* downstream region – forward primer, overlaps with *specR* |
| 810 | tcaatcttaccaaaacccacc | *lacA* downstream region – reverse primer |
| 805 | tggaggtgcggatgtgaaag | Nested forward primer for overlap extension PCR |
| 806 | ctcagccacttcaaagtcc | Nested reverse primer for overlap extension PCR |
| 803 | gatttcgattcctgttgccc | *lacA* deletion check (control) - forward primer for colony PCR |
| 10 | acattgttcagccgc | *lacA* deletion check (control) - reverse primer for colony PCR |
| 1015 | ggctacggtctcaggagccaaaccagccagctccgtatgc | *xylA* upstream region – forward primer with BsaI site |
| 18 | ctttgctgtttttcttctgctcaaagtagacgatcggc | *xylA* upstream region – reverse primer, overlaps with *xylA* down |
| 1016 | ggctacggtctccatggcaacagtaagtgaaacgtcagtcaccc | *xylA* downstream region – reverse primer with BsaI site |
| 19 | gccgatcgtctactttgagcagaagaaaaacagcaaag | *xylA* downstream region – forward primer, overlaps with *xylA* up |
| 776 | tttcgcagcaatgccaac | *xylA* deletion check (control) - reverse primer for colony PCR |
| 777 | cgccggatagacttgtttatac | *xylA* deletion check (control) – forward primer for colony PCR |
| pGGA FOR | ctgcaggaaggtttaaacgcatttagg | Plasmid sequencing primer |
| T7 FOR | gaaattaatacgactcactataggg | Plasmid sequencing primer |
| 1040 | gctacggtctcaggaggaggtgcggatgtgaaagcagc | *lacA* upstream – forward primer with BsaI site |
| 1041 | accaggtctcaccgttctcctactccttctcttcttgtcatttcagttgg | *lacA* upstream – reverse primer with BsaI site |
| 1042 | tggcggtctctacggttgtaaaagagatgaggctcattcgagg | *lacA* downstream – forward primer with BsaI site |

| | | |
|---|---|---|
| 1043 | ggctacggtctccatggcctcagccacttcaaagtccaagc | *lacA* downstream -reverse primer with BsaI site |
| 804 | ccgcttcaataagatgctctac | Reverse primer for *lacA* deletion check – colony PCR |
| 1017gga | gctacggtctcaaatgcgtaaaggagaagaacttttcactgg | *gfp* – forward primer with BsaI site |
| 1018gga | gctacggtctcctagattatttgtatagttcatccatgccatgtgtaatc | gfp – reverse primer with BsaI site |
| 1050 | acttggtctcacattctcctactccttctcttcttgtcatttcagttgg | *lacA* upstream region – reverse primer with BsaI site |
| 1051 | ggtctcatctattgtaaaagagatgaggctcattcgagg | *lacA* downstream region – forward primer with BsaI site |
| 815 | ggaatgcgctatatccaaacagaagg | Sequencing across *lacA* locus |
| 816 | taggagttgcttgacttcttctttaagc | Sequencing across *lacA* locus |
| 1072 | tagaggtctctagtcattgcaagccgttgcccacg | *lacA* downstream region – forward primer introducing stop codon (A replaces C) with BsaI site |
| 1073 | aggtctctgactataactcatgcatgcctggcttcttgg | *lacA* upstream region – reverse primer introducing stop codon (T replaces G) with BsaI site |
| 1052 | tgaaccgcgccaagaag | qPCR – forward primer for recombinant *lacA* |
| 1053 | gcaacggcttgcaatgact | qPCR – reverse primer for recombinant *lacA* |
| 1054 | ggaagtgctccgtgtcgaa | qPCR – forward primer for WT *lacA* |
| 1055 | tgtgacgctgtccatgcaa | qPCR – reverse primer for WT *lacA* |
| 1088 | gctacggtctcaggagttgatgggaatcacgagacag | *amyE* upstream region – forward primer with BsaI site |
| 1089 | accaggtctcaccgtcattcttgacactccttatt | *amyE* upstream region – reverse primer with BsaI site |
| 1090 | tggcggtctctacggtgagggcaaggctagacggg | *amyE* downstream region – forward primer with BsaI site |
| 1091 | ggctacggtctccatggatcggagtgagcgccacaagtg | *amyE* downstream region – reverse primer with BsaI site |
| 1097 | accaggtctctagacatggatgagcgatgatgatatc | *amyE* downstream region – forward primer introducing stop codon (A replaces C) with BsaI site |
| 1098 | tggtggtctctgtctactcttcatcatcattggcatac | *amyE* upstream region – reverse primer introducing stop codon (T replaces G) with BsaI site |
| 732 | cagtgccgtttaccgttcgcca | *amyE* deletion check (control) - forward primer for colony |
| 733 | atcggagtgagcgccacaagtg | *amyE* deletion check (control) - reverse primer for colony PCR |

(1) Koo BM, Kritikos G, Farelli JD, Todor H, Tong K, Kimsey H, et al. Construction and Analysis of Two Genome-Scale Deletion Libraries for Bacillus subtilis. Cell Syst. 2017 Mar;4(3):291-305.e7.

### Table S2. Sequences of homology arms

| | |
|---|---|
| *xylA* upstream | ccaaaccagccagctccgtatgcagccagcatcgccgcacccatgcctggcccttgctcgcttgtctgcttcacaattggcgt ttggaaaatgtcagcttgcatttgtagccacgtttcgctttttactcctccgccaatggaaacaattctctcaatcgtctttccttgttc tctgaacaaatcaagtgactctcttaatgaaaaagtgatgccttcaaggatcgacctagtaaagtcggccttcgtatggctgct atcagcaccgataaaactagcacggatagtagcgtcagcgtgaggcgtccgctctccggcaagatagggcgtaaacaa caacccatttgcccctgggggggacttctgccactcctgcaagcaattgcggaaacgtttcttcaggagcaaatgtttgcttaaa ccaacttaaactatatcctgccgctaatgtgacgcccattgtataaaacgcatcttcagctgcatggttaaaataatggacgc ctcctccaaatgtcttcccttttcttcctggtacgaaagcataacacctgacgtgccaatgctagcgagcgtcttgccgtttcta aaattccagctcctactgctccgcacgcattatcggctcctccagcaaaaacgggtgtttgcttggcagccccgtcgctaatg cgatttcttccgtcaactggccgacacaagcgtgtgattccactagcgtcggacaaagcgatggatcaatgccaaatgccg aacatagcggttcgctccatgtttttctttcgatatcaagcaagagtgtacctgctgcgtcagaatagtcggtatgcaatgcacc agtcaacttcaagcgcaaatagtcttttggaagcacgaatttggccgcttttgtaaacacttccggttcatgctcccgtacccac atcagctttggcaacgtaaaaccttcaagcgctggatttttggctaattgttgcagcctttcagcgcctagcttttcgtaaatcgct ttgcattgcggggaggtgcgtgtatcattccataaaatcgccggacgcaacacttcgttttgttcatcgagcaagacgagccc atgcatttgccccgaaaaactgattccctcaatttgagtcgttgcttgtggcaaggctgctgctatgtccgataatccagcaac ggtttccctgacccaatcgtttggatcttgctcgctccatcctgatttagggtgagacaacgggtatggtttagacacttccatga ccacttcacccgtctcgttaacgacaagcaatttcactgcgcttgtgcctaaatcaacaccaatgacatgtttcaatttgaatcg ctccttgccgatcgtctacttt |
| *xylA* downstream | gagcagaagaaaaacagcaaagaaaatgggaatgcataggacgtctgcattccctcgtaaactatgtttttagctcgttcct gcctcgctctcctgtgcgttgatcctttcatttcagcacaatttggtaccgttccattttttcgctaatgtctgtaatcgcttcccgtaat agcggcaagcaggagtcaaaacgctcctcattaaagcgataggctgggccggatatagtaatggaaccaacgatccgc ccttcatggaaaagcggcatcgccaaggcaatgacatcttccgtatattcgccatggctaatagcaaacccgcgtttgcgaa tggttgccagctcattacgcaattcaatttcacttgtcatcgtcctgtctgtataggctttcaaacctcccgcaatcacttcctcaat tgcctcgtcaggcatatacgccaagatgctgcggtaagacgcgcctacatataagggagcacggctgccagcagaaac ggaaaacttcactttattggctggctcaaccacttcaagcgtaagcgcttctttgccgtccagaattgttagaaacgtggactc gcctgtttttccgttaaacgggcaaggcttgggcggatcaattcttttacatttaactggtcgtacataatcgaacctagctccc acacggcaaacccgaggctatatttctttgtctctgtatctttcgttagaaagttttgggcttccagtgtttcaagtatccgataaatt ttcgtatgattgatccccgtagcgttggacagctccctgcctccccaaacaggcttttcttcgtaaacatttttaaaatctccaat gcttgatgtacagtttttaacacagctcctcctaggccattgcggcatacttaatcattactaagtttacatcaaaacactgcgc atgggtgactgacgtttcacttact |
| *lacA* upstream | tggaggtgcggatgtgaaagcagcggttacagttgaaccaattgctggattggatgatcgctttattaaaggagccgatatct cgatgttgccagaaatcgaggaaaacggcgggcgttattatgccgatggagaacaaattgatccactcgtattgttgcagg aaaagggcgtaaatgctgtccgtatccgcatttgggtagatccttacgatgaaaacggcaacccgtatggcggagggact gttgatggacagcgcgccgttgatctggcaaagcgcgcccatagctatggttttcaactgttgctcgattttcactacagtgattt ttggacagacccagggaagcagtttaagccaaaaagctgggaacattacagttttccggaattaaaggacgctatctatca ccatacatcaaatgtgcttggccagttaaaggatgcagggatcattccggaaatggtccaagttggcaatgaaattaatgcc ggaatgctatggcctgacggcaaaagctggggagaaggcggcggtgaatttgataggctcgccgcattgttgaaagcag ggcttactgccgtgaaagactgcgacccgggtatccgtacgatgcttcatttagcagaaggcggagatatcgatatggctca atggtggctcgatgagatcgggaagcgggacgttgattttgatgtcgttgggttgtcttactatccttactgggatggggatttca gtaagctcaaacaagtaatggagcttgttactaacgattatggcaaaggcgtcaatgtcgtcgaaacagcatatggctttac gacagacaatggcgacaacttagataatatcttttcatgtgaatacgcacagctagtcggttatccagcatcaccgcaaggg caagcgtcttacttgcgcgatttaatggcgacgattcacacggcaggaggagaagggttctattattgggaaccactttggat tccagtgtctggcgcctcttgggcgaacgaagccggaatgcgctatatccaaacagaaggggaagttggcaatgcttggg acaatcaagcgatgtttaattttaatggagaagcattgccttcattagatgtgttccatttagtaaaccaactgaaatgacaaga agagaaggagtaggaga |
| *lacA* downstream | ttgtaaaagagatgaggctcattcgaggattttgagcataataaatatgctgcactataacgaatataatgagctgcttcgagt attactagtaaaacaacacataagaccgatagccctacacggctgccggtcttcctttttatacaggtagaaataagaggg aaagaaattatttgatgttgggaaatatccaattaacttacattatcggaaaacatcgtctatactagtagtattatatattgatag aggtgatacgatgtcattaatgactgctgaaaaagctggggcgaaaattgttgagtggtatagttgtgtaattgctaagtctcat gatcaagctatttgcttaaagaagaagtcaagcaactcctaagtgagatgaaagacaatgataagatattggcctactattc tttagttgaatttaggcatgacatgctgataaatagatacaacaaaaatgaattacaggctgattttcagaatattgaacacat agctaaaattgacaacatgttaaagtacctctattactttgttagtggtcaaagcgagtatgtcaatgaacgctatagatcagct attaagctttttaataaagctcagcgcttgttagagtatgtgaacgatgaggctgaagaagctgagttttatcaatatagtgggc tagtttattacagattaaaccaataccttgtagctgcctctcatattgaacatgcaaaggttatctttgatcgtcttgaatatactga gccattattaaattgccaaattgtattagcaggtatttatcaagagttgcataaccctaaaaaagcagaggacatcttactaga tgccttggaaaaggccacggataatgaggtaatgctggggttaattaatcgatccttagggctgaataaacttggcactaaa gattacaaccaggcagaattttactttagacaagctttgtcttttaaggtgcacaaggatgcagcggttggagctaaaacaac atataatcttagtaatgttcttttcaaccaaggtaaccatgatgaggcaagaaagcagtttaaagcggctcatgcaggcgcta aatactacaaaaatcatgaatacatggcaagatgtctggcaacagaaggattgcatatcaaaaaagattacagtttagttg atacagcaatcgatgacctcaacaaattaggcttggactttgaagtggctgagg |

| | |
|---|---|
| *amyE* upstream | ttgatgggaatcacgagacaggttgcggttgaagcggttatgcttggagaaggggtcgtcaactgtgtgaacccgacatcc<br>ggcgttctcatggcggtgcttgccgccagcggtattccgtatgtcaagtggctgcggtttatggtgccgcttgctctgatttggttc<br>ttgatcgggcttgtctttatcgtgatcggagtcatgatcaattgggggccgttttaacgattgctgcccgccggcttgtacggcgg<br>gcttttgagttattcattgcagaagcgcaggctgttattgtaacatgtaagccataagccattcgtaaaagtgcgggaggaag<br>gtcatgaataatctgcgtaatagactttcaggcgtgaatgggaaaaataagagagtaaaagaaaaagaacaaaaaatct<br>ggtcggagattgggatgatagcgggagcatttgcgctgcttgatgtgatcatccgcggcattatgtttgaatttccgtttaaaga<br>atggctgcaagccttgtgtttttgttcatcattatcttatattactgcatcagggctgcggcatccggaatgctcatgccgagaat<br>agacaccaaagaagaactgcaaaaacgggtgaagcagcagcgaatagaatcaattgcggtcgcctttgcggtagtggt<br>gcttacgatgtacgacagggggattccccatacattcttcgcttggctgaaaatgattcttctttttatcgtctgcggcggcgttct<br>gtttctgcttcggtatgtgattgtgaagctggcttacagaagagcggtaaaagaagaaataaaaaagaaatcatctttttgttt<br>ggaaagcgagggaagcgttcacagtttcgggcagctttttttataggaacattgatttgtattcactctgccaagttgttttgatag<br>agtgattgtgataattttaaatgtaagcgttaacaaaattctccagtcttcacatcggtttgaaaggaggaagcggaagaatg<br>aagtaagagggatttttgactccgaagtaagtcttcaaaaaatcaaataaggagtgtcaagaatg |
| *amyE* downstream | tgagggcaaggctagacgggacttaccgaaagaaaccatcaatgatggtttctttttgttcataaatcagacaaaacttttct<br>cttgcaaaagtttgtgaagtgttgcacaatataaatgtgaaatacttcacaaacaaaaagacatcaaagagaaacataccc<br>tggaaggatgattaatgatgaacaaacatgtaaataaagtagctttaatcggagcgggttttgttggaagcagttatgcatttg<br>cgttaattaaccaaggaatcacagatgagcttgtggtcattgatgtaaataaagaaaaagcaatgggcgatgtgatggattt<br>aaaccacggaaaggcgtttgcgccacaaccggtcaaaacatcttacggaacatatgaagactgcaaggatgctgatattg<br>tctgcatttgcgccggagcaaaccaaaaacctggtgagacacgccttgaattagtagaaaagaacttgaagattttcaaag<br>gcatcgttagtgaagtcatggcgagcggatttgacggcattttcttagtcgcgacaaatccggttgatatcctgacttacgcaa<br>catggaaattcagcggcctgccaaaagagcgggtgattggaagcggcacaacacttgattctgcgagattccgtttcatgct<br>gagcgaatactttggcgcagcgcctcaaaacgtacacgcgcatattatcggagagcacggcgacacagagcttcctgttt<br>ggagccacgcgaatgtcggcggtgtgccggtcagtgaactcgttgagaaaaacgatgcgtacaaacaagaggagctgg<br>accaaattgtagatgatgtgaaaaacgcagcttaccatatcattgagaaaaaaggcgcgacttattatggggttgcgatga<br>gtcttgctcgcattacaaaagccattcttcataatgaaaacagcatattaactgtcagcacatatttggacgggcaatacggtg<br>cagatgacgtgtacatcggtgtgccggctgtcgtgaatcgcggagggatcgcaggtatcactgagctgaacttaaatgaga<br>aagaaaaagaacagttccttcacagcgccggcgtccttaaaaacattttaaaacctcattttgcagaacaaaaagtcaact<br>aaccgcaactttagagtaaagggctgattgtcaatgtgggagcagttgtatgatccgtttggaaacgagtatgtgagcgcact<br>tgtggcgctcactccgat |

**Table S3. Growth of *S. clausii* DSM 8716 in the presence of various antibiotics.** Sign (+) indicates visible growth, denoting resistance to the antibiotic; sign (–) indicates no growth, denoting sensitivity to the antibiotic. Highlighted boxes indicate antibiotic concentrations commonly used for selection in *B. subtilis*.

| | 1 µg $mL^{-1}$ | 5 µg $mL^{-1}$ | 10 µg $mL^{-1}$ | 20 µg $mL^{-1}$ | 50 µg $mL^{-1}$ | 100 µg $mL^{-1}$ |
|---|---|---|---|---|---|---|
| Ampicillin | + | + | + | + | + | + |
| Erythromycin | + | + | + | + | + | + |
| Nourseothricin | - | - | - | - | - | - |
| Spectinomycin | + | + | + | + | - | - |
| Kanamycin | + | + | + | + | + | - |
| Lincomycin | + | + | + | + | + | + |
| Tetracycline | + | - | - | - | - | - |
| Chloramphenicol | + | + | + | - | - | - |

**Figure S1.** Alignment of amino acid sequences of the xylose isomerase (XylA) from *B. subtilis* 168 and its homologue (XylA) in *S. clausii* DSM 8716. The multiple sequence alignment was generated using ClustalW as implemented in JALVIEW.

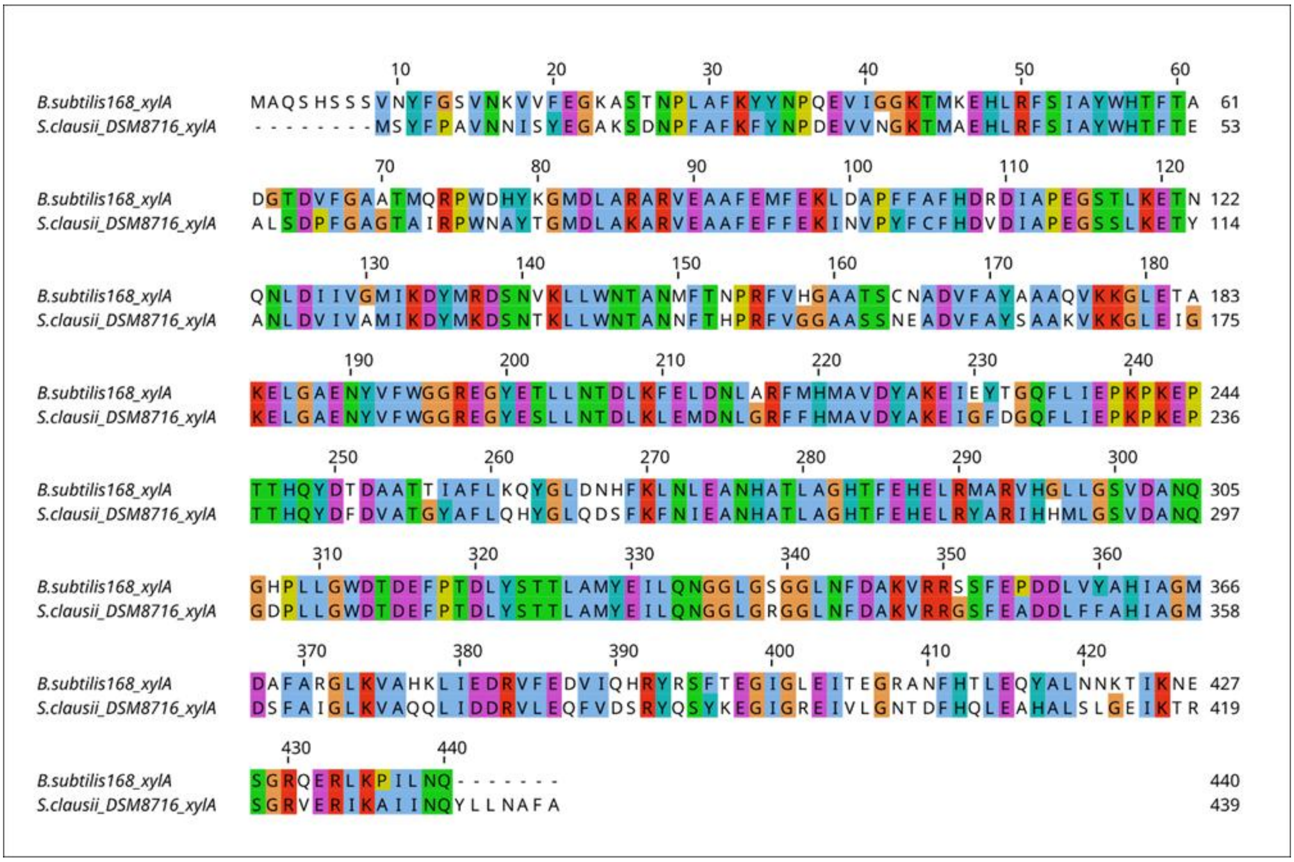

**Figure S2.** Alignment of amino acid sequences of the β-galactosidase GanA from *B. subtilis* 168 and its homologues (LacA) in *S. clausii* strains. The multiple sequence alignment was generated using ClustalW as implemented in JALVIEW.

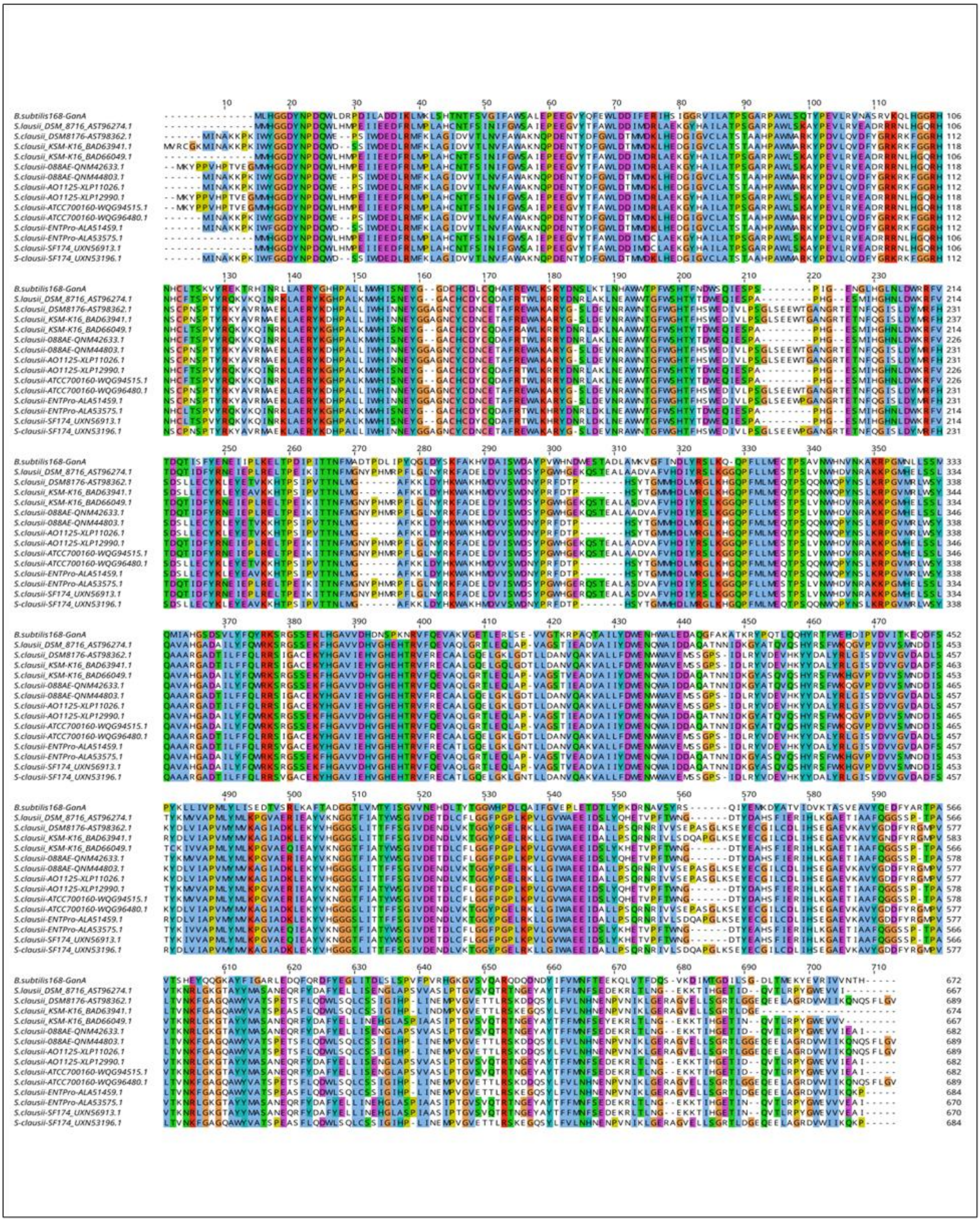

**Figure S3. Phylogenetic tree of** *B. subtilis* **β-galactosidases and various** *S. clausii* **strains.** The tree was constructed using sequences of the known GanA protein from *B. subtilis* 168 and the putative β-galactosidases from *S. clausii*, as described in the Materials and Methods section. Protein names are shown at the terminal nodes. The homologous LacA of *S. clausii* DSM 8716 is highlighted in red. The green box indicates the clade of putative LacA proteins.

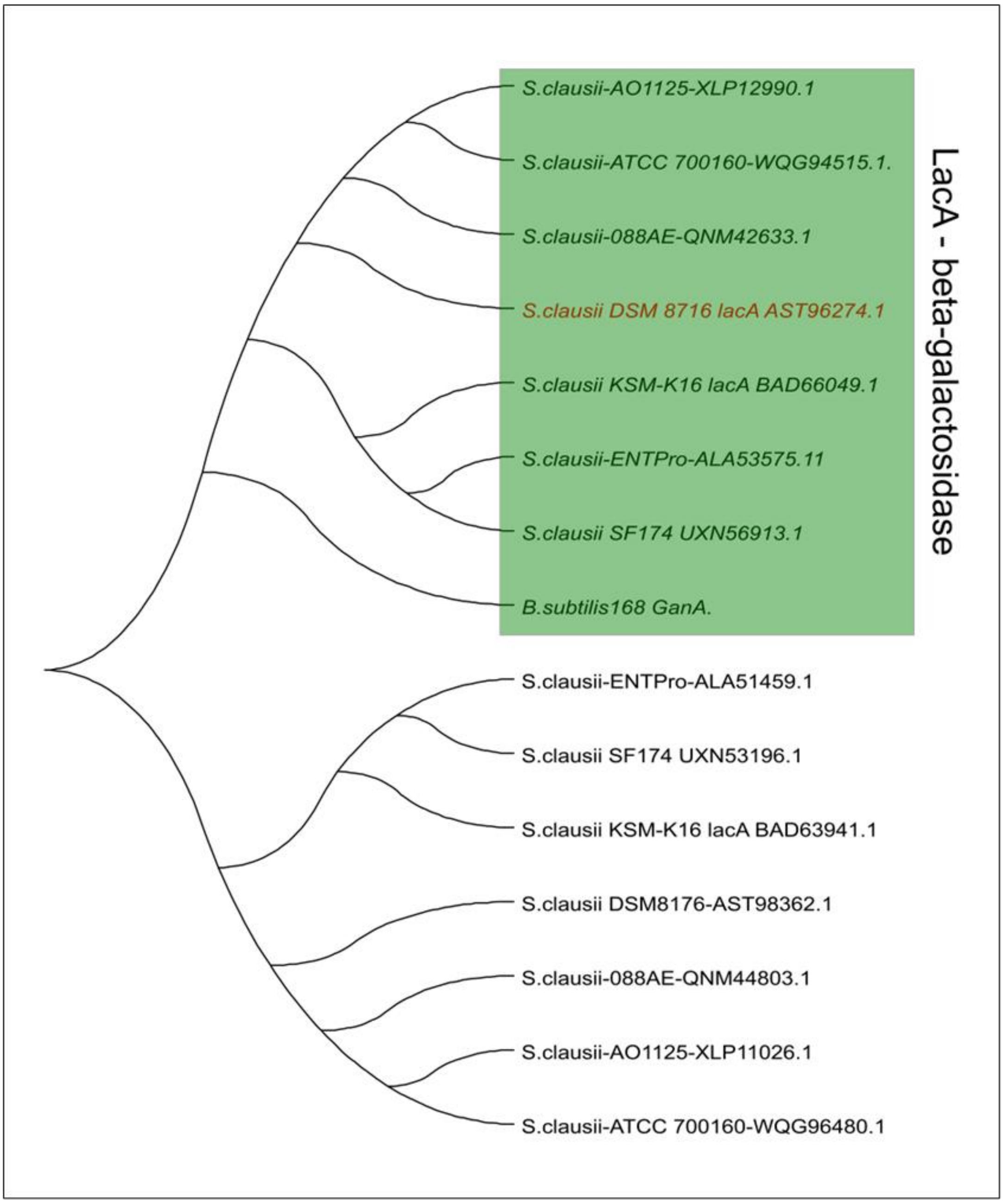

### Annex 1. Sequence of pM4B522

TCATGACCAAAATCCCTTAACGTGAGTTTTCGTTCCACTGAGCGTCAGACCCCGTAGAAAAGAT
CAAAGGATCTTCTTGAGATCCTTTTTTTCTGCGCGTAATCTGCTGCTTGCAAACAAAAAAACCA
CCGCTACCAGCGGTGGTTTGTTTGCCGGATCAAGAGCTACCAACTCTTTTTCCGAAGGTAACT
GGCTTCAGCAGAGCGCAGATACCAAATACTGTTCTTCTAGTGTAGCCGTAGTTAGGCCACCACT
TCAAGAACTCTGTAGCACCGCCTACATACCTCGCTCTGCTAATCCTGTTACCAGTGGCTGCTGC
CAGTGGCGATAAGTCGTGTCTTACCGGGTTGGACTCAAGACGATAGTTACCGGATAAGGCGCA
GCGGTCGGGCTGAACGGGGGGTTCGTGCACACAGCCCAGCTTGGAGCGAACGACCTACACC
GAACTGAGATACCTACAGCGTGAGCTATGAGAAAGCGCCACGCTTCCCGAAGGGAGAAAGGC
GGACAGGTATCCGGTAAGCGGCAGGGTCGGAACAGGAGAGCGCACGAGGGAGCTTCCAGG
GGGAAACGCCTGGTATCTTTATAGTCCTGTCGGGTTTCGCCACCTCTGACTTGAGCGTCGATT
TTTGTGATGCTCGTCAGGGGGGCGGAGCCTATGGAAAAACGCCAGCAACGCGGCCTTTTTAC
GGTTCCTGGCCTTTTGCTGGCCTTTTGCTCACATGTTCTTTCCTGCGTTATCCCCTGATTCTGT
GGATAACCGTCTGCAGGAAGGTTTAAACGCATTTAGGTGACACTATAGAAGTGTGTATCGCTCG
AGGGATCCGAATTCGAAGTCTTGGTACGGAGCGAGACCGGAGCGAGACGGGAGTCACGCAG
GTGTTGACGGCTAGCTCAGTGCTAGGTATTGTGCTAGCTTGACCCATTGTTTCAGCCAGGAGG
TTCTATGTCAGTAATCGCTAAGCAGATGACATACAAGGTTTACATGTCTGGTACAGTGAACGGA
CACTACTTTGAAGTAGAGGGCGATGGTAAAGGGAAGCCTTATGAAGGAGAGCAGACGGTCAA
GTTAACAGTCACCAAAGGCGGGCCTCTGCCATTCGCATGGGATATTTTAAGTCCTCAATGCCAG
TATGGATCTATACCATTCACTAAGTATCCAGAAGATATACCTGATTATGTGAAACAGAGTTTTCCC
GAAGGCTACACATGGGAACGCATAATGAATTTTGAGGACGGTGCTGTGTGCACAGTGAGTAAT
GATTCTTCTATACAGGGTAATTGCTTCATTTATCATGTAAAGTTCAGCGGTCTGAATTTTCCACCT
AATGGTCCAGTAATGCAGAAGAAAACCCAGGGTTGGGAGCCAAACACCGAGCGACTGTTCGC
ACGCGATGGTATGCTCTTAGGTAACAACTTCATGGCTCTGAAGTTAGAGGGCGGTGGGCATTA
CTTATGTGAATTTAAGACAACGTATAAAGCCAAGAAGCCGGTGAAGATGCCTGGTTACCACTAC
GTTGACAGAAAGTTGGATGTCACCAACCATAATAAGGACTATACGTCTGTTGAGCAATGCGAAA
TTAGTATAGCCAGAAAACCCGTCGTGGCATAATAATACTAGAGTCACACTGGCTCACCTTCGGG
TGGGCCTTTCTGCGTTTATATACTAGAGAGAGAATATAAAAAGCCAGATTATTAATCCGGCTTTTT
TATTATTTCACCTGCAGGACCATCGTCTCACCATGGTCTCACCATTCCTGTAGACTTCTTAATTA
AGACGTCAGAATTCTCGAGGCGGCCGCATGTGAGTCTCCCTATAGTGAGTCGTATTAATTTCGG
TCTAGTTAATGTGTAACGTAACATTAGCTAGATTTTTTTATTCAAAAAAATATTTACAAATATTAGGA
AATTTAAGTGTAAAAGAGTTGATAAATGATTATATTGGGACTATAATATAATTAAGGATCTGGCTTC
CTCCGAAGACGTTATCAAAGAGTTCATGCGTTTCAAAGTTCGTATGGAAGGTTCCGTTAACGGT
CACGAGTTCGAAATCGAAGGTGAAGGTGAAGGTCGTCCGTACGAAGGTACCCAGACCGCTAA
ACTGAAAGTTACCAAAGGTGGTCCGCTGCCGTTCGCTTGGGACATCCTGTCCCCGCAGTTCC
AGTACGGTTCCAAAGCTTACGTTAAACACCCGGCTGACATCCCGGACTACCTGAAACTGTCCT
TCCCGGAAGGTTTCAAATGGGAACGTGTTATGAACTTCGAAGACGGTGGTGTTGTTACCGTTA
CCCAGGACTCCTCCCTGCAAGACGGTGAGTTCATCTACAAAGTTAAACTGCGTGGTACCAACT
TCCCGTCCGATGGTCCGGTTATGCAGAAAAAAACCATGGGTTGGGAAGCTTCCACCGAACGTA
TGTACCCGGAAGACGGTGCTCTGAAAGGTGAAATCAAAATGCGTCTGAAACTGAAAGACGGT
GGTCACTACGACGCTGAAGTTAAAACCACCTACATGGCTAAAAAACCGGTTCAGCTGCCGGGT
GCTTACAAAACCGACATCAAACTGGACATCACCTCCCACAACGAAGACTACACCATCGTTGAA
CAGTACGAACGTGCTGAAGGTCGTCACTCCACCGGTGCTTAATAAAGGCCTCGATCCCGCAAG
AGGCCCGGCAGTACCGGCATAACCAAGCCTATGCCTACAGCATCCAGGGTGACGGTGCCGAG
GATGACGATGAGCGCATTGTTAGATTTCATACACGGTGCCTGACTGCGTTAGCAATTTAACTGT
GATAAACTACCGCATTAAAGCTTATCGATTATGTCTTTTGCGCAGTCGGCTTAAACCAGTTTTCG
CTGGTGCGAAAAAAGAGTGTCTTGTGACACCTAAATTCAAAATCTATCGGTCAGATTTATACCGA
TTTGATTTTATATATTCTTGAATAACATACGCCGAGTTATCACATAAAAGCGGGAACCAATCATCA
AATTTAAACTTCATTGCATAATCCATTAAACTCTTAAATTCTACGATTCCTTGTTCATCAATAAACT

CAATCATTTCTTTAATTAATTTATATCTATCTGTTGTTGTTTTCTTTAATAATTCATCAACATCTACACCGCCATAAACTATCATATCTTCTTTTTGATATTTAAATTTATTAGGATCGTCCATGTGAAGCATATATCTCACAAGACCTTTCACACTTCCTGCAATCTGCGGAATAGTCGCATTCAATTCTTCTGTTAATTATTTTTATCTGTTCATAAGATTTATTACCCTCATACATCACTAGAATATGATAATGCTCTTTTTTCATCCTACCTTCTGTATCAGTATCCCTATCATGTAATGGAGACACTACAAATTGAATGTGTAACTCTTTTAAATACTCTAACCACTCGGCTTTTGCTGATTCTGGATATAAAACAAATGTCCAATTACGTCCTCTTGAATTTTTCTTGTTTTCAGTTTCTTTTATTACATTTTCGCTCATGATATAATAACGGTGCTAATACACTTAACAAAATTTAGTCATAGATAGGCAGCATGCCAGTGCTGTCTATCTTTTTTTGTTTAAAATGCACCGTATTCCTCCTTTGCATATTTTTTTATTAGAATACCGGTTGCATCTGATTTGCTAATATTATATTTTTCTTTGATTCTATTTAATATCTCATTTTCTTCTGTTGTAAGTCTTAAAGTAACAGCAACTTTTTTCTCTTCTTTTCTATCTACAACTATCACTGTACCTCCCAACATCTGTTTTTTTCACTTTAACATAAAAAACAACCTTTTAACATTAAAAACCCAATATTTATTTATTTGTTTGGACAATGGACAATGGACACCTAGGGGGGAGGTCGTAGTACCCCCCTATGTTTTCTCCCCTAAATAACCCCAAAAATCTAAGAAAAAAAGACCTCAAAAAGGTCTTTAATTAACATCTCAAATTTCGCATTTATTCCAATTTCCTTTTTGCGTGTGATGCGCTGCGTCCATTAAAAATCCTAGAGCTTTGCAACCGAAAGTTAATAGCTGTCGCTACTACTTTCGCTTACGCTCTAAGTATATTTTAAGGACTGTCACACGCAAAAAGTTTTCTCGGCATAAAAGTACCTCTACATCTCTAAATCGTCTGTACGCTGTTTCTCACGCTTTCTATCGACCTTCTGGACATTATCCTGTACAACATCCATAAACTGTCCCACACGCTCAAATTTGGAATCATTAAAGAATTTCTCTTTAAGCCTATTAAACCCTTTCTCAAACCCAGGGAAATTCGCCCTCGCAGCACGATATAAAGTCACTGTACTAGCTTGAAATTTCTCTGATACATTCAACTGCTCATTCAAACTATCATTCTCTCGCTTTAATTTATTAACCTCTTTACTTTTTTCGTGATACCCCTCTTTCCATGTATTCACTACTTCTTTCAAACTCTCTCTACGTTTTTTTAATTCTTGATTTTCTGTGTAATAGTCTGTGCTCTTAATATTTTCGTAATCATCAACAATCCGTTCTGCAGAAGAGATTGTTTCTTGCAGGCGTTCAAATTCATCAGCAGTTAATATCTTTCTACCAGTCTCTTCACGTCCAGAGAACAAACCTGTACGCTCATTTTCATAATCAAAGGGTTTCGTAGACCTCATATGCTCTATTCCACTCTGTAACTGCTTATTTGCCTTCTGTAACTCATCCTTAACTTCTTGCAGTTCCTGTTTATGAAATACAGTATCTTTCTTGTACTGATCCATCGCTTTATGTTCTCGTTCTGTAACCTCTTTGGACGTGCCTCTTTCAAGTTCATAACCTTTCTCATTCACATACTCATTAAATCTATCTTGTAATTGAGTAAAGTCTTTCTTGTTGCCTAACTGTTCTTTTGCAGACAATCTCCCGTCCTCTGTTAAAGGGACAAAACCAAAGTGCATATGTGGGACTCTTTCATCCAGATGGACAGTCGCATACAGCATATTTTCCTTACCGTATTCATTTTCTAGAAACTCCAAGCTATCTTTAAAAAATCGTTCTATTTCTTCTCCGCTTAAATCATCAAAGAAATCTTTATCACTTGTAACCAGTCCGTCCACATGTCGAATTGCATCTGACCGAATTTTACGTTTCCCTGAATAATTCTCATCAATCGTTTCATCAATTTTATCTTTATACTTTATATTTTGTGCGTTAATCAAATCATAATTTTTATATGTTTCCTCATGATTTATGTCTTTATTATTATAGTTTTTATTCTCTCTTTGATTATGTCTTTGTATCCCGTTTGTATTACTTGATCCTTTAACTCTGGCAACCCTCAAAATTGAATGAGACATGCTACACCTCCGGATAATAAATATATATAAACGTATATAGATTTCATAAAGTCTAACACACTAGACTTATTTACTTCGTAATTAAGTCGTTAAACCGTGTGCTCTACGACCAAAACTATAAAACCTTTAAGAACTTTCTTTTTTTACAAGAAAAAAGAAATTAGATAAATCTCTCATATCTTTTATTCAATAATCGCATCCGATTGCAGTATAAATTTAACGATCACTCATCATGTTCATATTTATCAGAGCTCGTGCTATAATTATACTAATTTTATAAGGAGGAAAAAATATGGGCATTTTTAGTATTTTTGTAATCAGCACAGTTCATTATCAACCAAACAAAAAATAAGTGGTTATAATGAATCGTTAATAAGCAAAATTCATATAACCAAATTAAAGAGGGTTATAGAATTCTTGACATTTTTCTTGTGGATCTGTATAATAAAGAATAATTATTAATCTGTAGACAAATTGTGAAAGGATGTACTTAAACGCTAACGGTCAGCTTTATTGAACAGTAATTTAAGTATATGTCCAATCTAGGGTAAGTAAATTGAGTATCAATATAAACTTTATATGAACATAATCAACGAGGTGAAATCATGAGCAATTTGATTAACGGAAAAATACCAAATCAAGCGATTCAAACATTAAAAATCGTAAAAGATTTATTTGGAAGTTCAATAGTTGGAGTATATCTATTTGGTTCAGCAGTAAATGGTGGTTTACGCATTAACAGCGATGTAGATGTTCTAGTCGTCGTGAATCATAGTTTACCTCAATTAACTCGAAAAAAACTAACAGAAAGACTAATGACTATATCAGGAAAGATTGGAAATACGGATTCTGTTAGACCACTTGAAGTTACGGTTATAAATAGGAGTGAAGTTGTCCCTTGGCAATATCCTCCAAAAAGAGAATTTATATACGGTGAGTGGCTCAGGGGTGAATTTGAGAATGGACAAATTCAGGAACCAAGCTATGATCCTGATTTGGCTATTGTTTTAGCACAAGCAAGAAAGAATAGTATTTCTCTATTTGGTCCTGATTCTTCAAGTATACTTGTCTCCGTACCTTTGACAGATATTCGAAGAGCAATTAAGGATTCTTTGCCAGAACTAATTGAGGGGATAAAAGGTGATGAGC

GTAATGTAATTTTAACCCTAGCTCGAATGTGGCAAACAGTGACTACTGGTGAAATTACCTCGAAA
GATGTCGCTGCAGAATGGGCTATACCTCTTTTACCTAAAGAGCATGTAACTTTACTGGATATAGC
TAGAAAAGGCTATCGGGGAGAGTGTGATGATAAGTGGGAAGGACTATATTCAAAGGTGAAAGC
ACTCGTTAAGTATATGAAAAATTCTATAGAAACTTCTCTCAATTAAGGCTAATTTTATTGCAATAAC
AGGTGCTTACTTTTAAAACTACTGATTTATTGATAAATATTGAACAATTTTTGGGAAGAATAAAGC
GTCCTCTTGTGAAATTAGAGAACGCTTTATTACTTTAATTTAGTGAAACAATTTGTAACTATTGAT
CGATCTCGAG
